# Subsets of NLR genes drive adaptation of tomato to pathogens during colonisation of new habitats

**DOI:** 10.1101/210559

**Authors:** Remco Stam, Gustavo A. Silva-Arias, Aurelien Tellier

## Abstract

- Nucleotide binding site, Leucine-rich repeat Receptors (NLRs), are canonical resistance (R) genes in plants, fungi and animals, functioning as central (helper) and peripheral (sensor) genes in a signalling network. We investigate NLR evolution during the colonisation of novel habitats in a model tomato species, *Solanum chilense.*
- We used R-gene enrichment sequencing (RENSeq) to obtain polymorphism data at NLRs of 140 plants sampled across 14 populations covering the whole species range. We inferred the past demographic history of habitat colonisation by resequencing whole genomes from three *S. chilense* plants from three key populations, and performing Approximate Bayesian Computation using data from the 14 populations.
- Using these parameters we simulated the genetic differentiation statistics distribution expected under neutral NLR evolution, and identified small subsets of outlier NLRs exhibiting signatures of selection across populations.
- NLRs under selection between habitats are more often helper genes, while those showing signatures of adaptation in single populations are more often sensor-NLRs. Thus, centrality in the NLR network does not constrain NLR evolvability, and new mutations in central genes in the network are key for R gene adaptation during colonisation of different habitats.

## INTRODUCTION

Antagonistic interactions can generate endless coevolution between hosts and their pathogens. The Red Queen hypothesis predicts that the genomes of both interacting partners evolve to match each other’s changes (Van Valen, 1973): pathogens evolve infectivity to overcome defences, while hosts evolve pathogen recognition and resistance to avoid infection. Changes in allele frequencies of different infectivity/resistance specificities occur over tens to thousands of generations at the loci determining the outcome of interaction. Two extreme types of dynamics have been proposed, which differ in their signatures at the phenotypic (Gandon *et al.*, 2008) and genotypic polymorphism levels (Woolhouse *et al.*, 2002): the arms race model (Bergelson *et al.*, 2001) is characterised by recurrent selective sweeps in both partners, and the trench warfare model (Stahl *et al.*, 1999) shows long-lasting balancing selection. These premises form the basis for genome scans to detect the genes under coevolution/selection in hosts and pathogens (Bakker *et al.*, 2006). Plant species exhibit a spatial distribution across habitats, which influences coevolutionary dynamics (Gandon *et al.*, 2008; Parratt *et al.*, 2016). Diverse habitats generate differential pathogen pressure across space due to variation in 1) disease presence or absence and prevalence, 2) disease transmission between hosts, and 3) co-infection or competition between pathogen species. As a result, spatial heterogeneity is observed for infectivity in pathogens and resistance in hosts (Thrall *et al.*, 2001; Caicedo & Schaal, 2004). Species expansion and colonisation of new habitats could in addition cause the host to encounter new pathogens and subsequently promote coevolutionary dynamics at single copy genes or gene families compared to the original habitat. Despite the wealth of studies at the phenotypic and ecological levels (Thrall & Burdon, 2003; Thrall *et al.*, 2012; Tack & Laine, 2014), we know little about the genetic basis of host-pathogen coevolution in spatially heterogeneous populations and during the colonisation of new habitats. A crucial issue for such studies is to disentangle the signatures of selection at a few genes from the genome-wide effect of demography in shaping diversity. The problem is especially difficult in searches of genes under selection (selective sweeps or balancing selection) during adaptation to new habitats, because colonisation events generate bottlenecks resulting in an increase of the variance of the measured nucleotide diversity over the genome (e.g. in *Arabidopsis thaliana* (Lee *et al.*, 2017; Exposito-Alonso *et al.*, 2018) and *Arabis alpina* (Laenen *et al.*, 2018)).

Resistance genes are the key players in host - pathogen interactions, as they are sensing pathogen molecules to activate immune responses. Canonical R genes are members of the NLR family NLR (nucleotide binding site, leucine-rich repeat containing receptor) that occurs in both plants and animals (Jones *et al.*, 2016). NLRs have a modular structure. NLRs can have a N-terminal TIR-domain (TNLs) or CC-domain (CNLs), followed by a Nucleotide Binding Site and leucine rich repeats). In *A. thaliana*, some NLRs appear to show signatures of positive or balancing selection (Bakker *et al.*, 2006) and overall NLRs seem to show more positive selection than other defence-related gene families (Mondragón-Palomino *et al.*, 2017). Yet, detailed studies of NLR evolution in wild pathosystems are lacking. In most cases only few candidate NLR genes have been studied. For example, in the common bean (*Phaseolus vulgaris*) the NLR locus PRLJI1 shows slightly higher overall *F*_*ST*_ and markedly different patterns of spatial differentiation within and between populations compared to the genome-wide average (AFLP markers) (De Meaux *et al.*, 2003). In wild emmer wheat (*Triticum dicoccum)* a marker-based analysis shows that NLRs exhibit higher differentiation (*F*_ST_ = 0.58) than other markers (*F*_ST_ = 0.38) (Sela *et al.*, 2009). Within a single genus the number of NLRs can differ dramatically between species suggesting that the NLR family experiences a rapid birth-and-death process (Michelmore & Meyers, 1998) driven by large scale gene duplication and deletion, whereas within species variation is hypothesised to be mainly found at the nucleotide level at a few key genes (Wu *et al.*, 2017b) or at a few duplicated genes (Hörger *et al.*, 2012). The evolutionary mechanism explaining the latter is termed as the recycling of existing NLRs (Holub, 2001) by generation of new specificity at a given locus entering the host-pathogen coevolutionary process.

These theoretical expectations are based on the evolution of NLRs as single genes “sensing” the presence of pathogens (either directly or indirectly, Kourelis & Hoorn, 2018). It has now been found that NLRs form a complex multi-layer signalling network (Wu *et al.*, 2018) to recognise pathogens and transduct the signal into the appropriate defence response. A major recent finding is that members of the NRC (NLR required for cell death) clade are for example central in the network and are required as “helpers” for the functioning of other, “sensor” NLRs (Wu *et al.*, 2017a, 2018). The sensor NLRs are more peripheral in the network and have less connectivity to other genes. Expanding on the previous questions, we want to investigate if all NLRs in the network have the same evolutionary potential when colonising new habitats and encountering new pathogens.

We designed our study to address the following questions. How many NLR genes are involved in coevolution with pathogens across populations? What is the time scale of coevolution in newly colonised habitats and which genes are involved? We are particularly interested in finding how many genes exhibit different selection pressures between the original and the newly colonised habitat, namely genes evolving neutrally in the original habitat and being under (positive or balancing) selection in the derived one. Finally, we also want to know whether there are differences in evolutionary changes for the various annotated NLR classes. For example, are genes central in the network showing more evolutionary constraints?

We answer these questions by studying the sequence evolution of NLR genes in a wild tomato species, *Solanum chilense*. This species is particularly amenable to this approach: it exhibits a high effective population size (*N*_e_), high nucleotide diversity (heterozygosity), and high recombination rates (Arunyawat *et al.*, 2007). These features are due to outcrossing, spatial structuring of populations linked by gene flow and the presence of seed banks (Arunyawat *et al.*, 2007; Tellier *et al.*, 2011). *S. chilense* occurs in southern Peru and northern Chile. Local adaptation to abiotic and biotic stresses in *S. chilense* or its sister species is indicated by 1) signatures of positive selection in genes involved in cold and drought stress response (Xia *et al.*, 2010; Fischer *et al.*, 2013; Nosenko *et al.*, 2016; Böndel *et al.*, 2018), 2) balancing selection in several genes of the *Pto* resistance pathway providing resistance to *Pseudomonas sp*. (Rose *et al.*, 2011), and 3) variable resistant phenotypes against filamentous pathogens across populations (Stam *et al.*, 2017). *S. chilense* is also an established source of fungal and viral R genes used in breeding programmes (Tabaeizadeh *et al.*, 1999; Verlaan *et al.*, 2013). *S. chilense* consists of four clearly defined geographical groups. The central group, considered the centre of origin of the species, is found in the mesic part of its range in southern Peru and northern Chile. Two southern groups likely result from two distinct southward colonisation events around the Atacama desert, one towards the coastal part of northern Chile (southern coast group), and other through high altitudes of the Chilean Andes (southern mountain group) (Böndel *et al.*, 2015). The northern group (southern Peru) was derived from the central one and is found in sympatry with its sister species *S. peruvianum*. The bottlenecks during these colonisation events have been relatively mild, so that this species still exhibits high genetic diversity (and adaptive potential) after the range expansions. The southward colonisation events provide two independent replicates of the process of adaptation to new abiotic and biotic stresses. In a recent study, we sequenced the ∼915 Mb reference genome and *de novo* transcriptome of *S. chilense* (Stam *et al.*, 2019). We annotated 25,885 high confidence gene models, 71% of them are supported by transcriptome data. Our annotation yielded 236 NLRs in *S. chilense*, 201 can be considered high quality annotations, and all previously identified NLR functional clades (Jupe *et al.*, 2013; Andolfo *et al.*, 2014) can be found, albeit some with different numbers compared to other tomato species. Additionally, we identified two newly expanded clades. Overal, the *S. chilense* NLR complement looked similar to that of *S. pennellii*, a wild tomato species for which we have previously shown that NLR sequence diversity is maintained within a single population (Stam *et al.*, 2016).

We derive a three-pronged approach to examine the adaptation of *S. chilense* NLR genes between populations of different habitats. We re-sequence all NLRs in 14 populations for ten plants per population. Then, we infer the colonisation and demographic history based on three full genomes representative of the three major habitat groups [central (centre of origin), southern coastal and southern mountain (derived)] and use these data to infer expected NLR diversity in all fourteen populations. Lastly we combine these data to identify NLRs under different selection in the derived groups compared to the original one. We conclude by discussing the selective pressures acting on and the evolvability of the host defence network when colonising new habitats in the light of the functional classes to which the NLRs belong.

## Methods

### Plant material and accessions

We grew ten plants for each of the 14 populations of *S. chilense* in our glasshouse (20°C, 16h light). Accession numbers: LA3111, LA4330, LA2932, LA1958, LA1963, LA2747, LA2755, LA2931, LA3784, LA3786, LA2750, LA4107, LA4117(A), LA4118. (Supplementary Notes, S1)

### Pooled R gene enrichment sequencing and SNP analysis (RENSeq)

Genomic DNA was extracted from ten mature plants per population and pooled. Sequencing was done at NGS@TUM. We performed the library preparation, read mapping and SNP calling as described before (Stam *et al.*, 2016) and (Supplementary Notes S2). The NLR probes were based on known R-genes in solanaceae and *A. thaliana* and have successfully been used before (Stam *et al.*, 2016). Mapping was done using Stampy (Lunter & Goodson, 2011), SNP calling using two callers: GATK (McKenna *et al.*, 2010) and Popoolation (Kofler *et al.*, 2011). We previously found 236 NLRs in *S. chilense* and focus here on 201 high quality ones (Stam *et al., 2019)*. To verify the stringency of the filters and the cut-off values, we compared the merged SNP calls to Sanger sequence data for three genes for all ten plants for several populations (Supplementary data 3). After comparison, cut-offs were adjusted to obtain the best true SNP calls and both callers were run again. The combined results of the last round were used. Summary statistics π, θ _W_, π_N_ and π_S_ were calculated with SNPGenie (Nelson *et al.*, 2015) *F*_ST_ values were calculated for pairs of populations using the Hudson et al. (Hudson *et al.*, 1992) estimator: *F*_ST_ = (π_between -_ π_within_)/ π_between_. We assure robustness of the *F*_ST_ calculations by using only 91 NLR genes with high and even coverage between all compared populations. Significant differences were tested using ANOVA, with the Tukey HSD test and recorded when p < 10^−5^, unless stated otherwise.

In addition, we sequenced 14 reference loci (hereafter CT loci), used in previous studies in *S. chilense* and *S. peruvianum* (e.g. Arunyawat *et al.*, 2007; Böndel *et al.*, 2015). The CT loci summary statistics were compared to the results by Böndel et al. (2015) who used an overlapping set of populations (but not the same plants). Due to known difficulty to reliably assess allele frequencies in pooled data (Futschik & Schlotterer, 2010), our analyses are based on the nucleotide diversity statistics. These seem well estimated by our SNP call procedure when we compare at the CT loci our results to a previous study (see results below).

### Full Genome Resequencing

Accessions LA4330 and LA2932, representing southern mountain and southern coast, respectively, were sequenced at Eurofins Genomics on a Illumina HiSeq 2500 with standard library size of 300bp. We mapped the sequenced reads of the three sequenced plants (our reference genome (LA3111), representing the central region and resequence data from LA4300 and LA2932 representing the southern mountain and southern coast populations, respectively), against our *S. chilense* reference genome (Stam *et al.*, 2019) using BWA (mem, call -M with default parameters). SNPCalling was done using samtools (mpileup -q 20 -Q 20 -C 50).

### Demographic inferences with MSMC, ABC and simulation of summary statistics

We inferred the demographic history of three *S. chilense* populations LA3111, LA2932 and LA4330 using whole genome sequence data and the MSMC method (Schiffels & Durbin, 2014). MSMC relies on long genomic fragments, thus we restrict the variant calling to the 200 largest scaffolds of the *S. chilense* reference genome: ∼79.6Mb of sequence (mean length=398Kb, min=294Kb, max=1.12Mb). We estimate the past changes in effective population (*N*_e_) size per population and cross-coalescence rates, assuming a per site mutation rate of 5×10^−8^ and generation time of 5 years. (full details on data preparation and settings are given in Supplementary Notes S3). The latter rates compare the frequency at which the most recent common ancestor is found either within individual (diploid) genomes or between two individuals of different populations, and thus indicate the time of population split. To check robustness of the inference we simulated independent scenarios of demography and divergence using ms (Hudson, 2002). We tested the demographic estimation with simulated sequences of the same length as the *S. chilense* reference genome and same estimated values of the population mutation rate (based on θ_W_ values) and the population recombination rate (based on ρ values). We assessed the ability of MSMC to estimate the correct demographic parameters (population sizes, time of split) for simple demographic models, and a model of population splits mimicking the southward colonisation events. Using the two simulated scenarios that better resembled the MSMC estimations obtained with the observed data, we simulated sequences with the same features as our empiric NLR dataset to obtain neutral distributions of the summary statistics (gene length =2149).

Given that the estimates obtained with MSMC do not assume migration between populations, a feature of many wild plant species which likely occurs between populations of *S. chilense* (Tellier *et al.*, 2011; Böndel *et al.*, 2015) we additionally implemented a more comprehensive demographic inference via an Approximate Bayesian Computation (ABC) approach (Beaumont *et al.*, 2002). This allows us to take into account post-split gene flow between populations and test for the most likely divergence scenario (Supplementary Notes S4). Three demographic models of geographic group divergence were tested to assess the order of the splits. We then estimated *N*_e_, divergence times and migration rates under the best supported model. The data used for the ABC consist of synonimous sites of the 91 high quality NLRs and 14 CT reference loci at the 14 populations. The ABC is conducted with ms (Hudson, 2002) and the R package abc (Csilléry et al., 2012). From the ABC posterior parameter estimations we generated a set of neutral distributions of *F*_ST_ values for all population pairwise comparisons which, based on 30,000 loci defined by the average length of our NLRs and genomic population recombination rate estimated with MSMC (4.5×10^−9^ – 1.1×10^−8^ per site per generation).

Using forward simulations (Supplementary Notes, S5) we tested that genes under different selective pressures in different populations can be revealed by outlier high *F*_ST_ values compared to the neutral expected distributions from our neutral demographic scenario. For that, we ran simulations using SLiM (Haller & Messer, 2019) assuming genes evolving neutrally in all populations, and changing from neutral to either positive or balancing selection in the southward colonisation processes.

### Definition of outlier NLR

For each pairwise comparison between the populations, we conservatively selected the NLRs that fell outside the maximum simulated value (out of 30,000 simulations). Main habitat adaptation genes were defined by selecting the genes that occur as outlier in at least one third of the possible pairwise population comparisons between two groups. To test whether the relative abundance of the NLR classes in the main and local adaptation groupings could arise by chance, we randomised the *F*_ST_ values within 1) the whole data set, and 2) the total set of selected outlier NLRs, and subsequently reran our analyses. Using these randomisation outputs we calculated the average number of major genes that can be identified (under 1,000 whole dataset randomisation) or the mean number of NRC genes that are classified as major genes (in 1,000 randomisations following procedure 2 within outliers). We estimated the confidence interval for the number of major genes to be found from the random sampling (mean ±2σ). (Supplementary Notes, S6)

## Results

### Enrichment sequencing provides high coverage and reliable summary statistics

Polymorphism data at NLRs were obtained by targeted enrichment sequencing of pooled DNA of ten plants for each of the 14 populations (Figure 1A). For each population one to two million read pairs passed trimming and quality controls (Supplemental Data 01). For all pooled samples, the coverage exceeds 100x for 80% of the targeted NLRs. To evaluate the short-read data quality, we also enriched and sequenced the set of 14 CT genes, which showed a coverage of more than 100x in most pooled samples (S Figure 1A). We called SNPs per gene against our LA3111 reference genome (Stam *et al.*, 2019) for the 201 high quality NLRs (out of 236 identified genes) and all 14 CT genes in each population. We calculated the statistic π, summarising nucleotide diversity, and π_N_ and π_S_ as the nucleotide diversity for non-synonymous or synonymous sites only (Supplemental Data 2). No significant correlation was found between the number of mapped reads or bases and the number of SNPs per population (R^2^ = 0.46 and p = 0.1) or π per population (for read pairs: R^2^ = 0.30, p = 0.30, for bases: 0.35 and 0.2) (S Figure 1B). Thus, our data is not biased for coverage differences between the samples.

**Figure 1.**
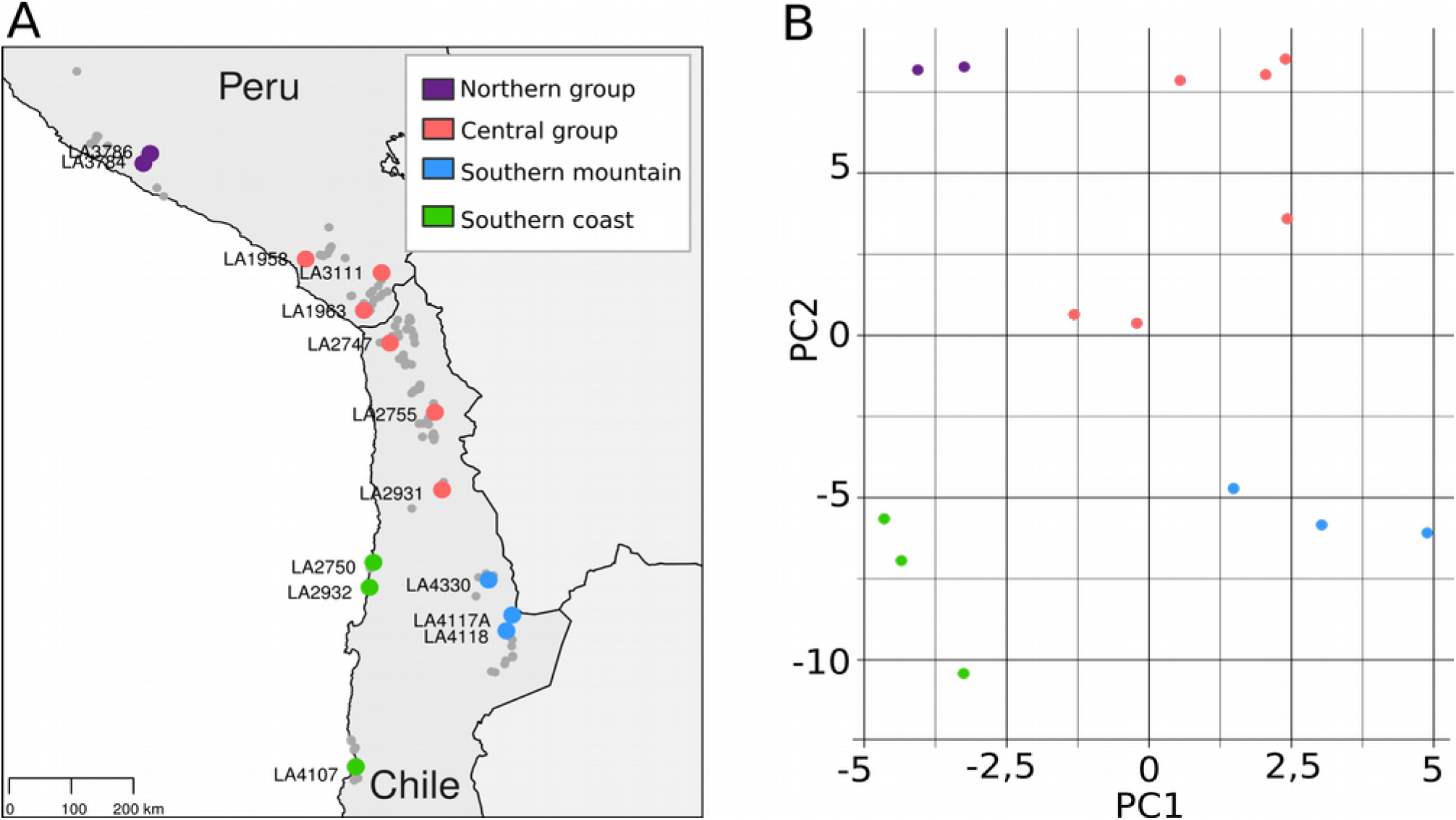
Overview of the studied populations and structuring of species-wide NLR diversity across the 14 populations. A) Map of the studied populations (colored by genotype group) compared to all reported *S. chilense* populations from the TGRC database, UC Davis, USA (grey dots). B) Principal component analysis of all SNPs in all sequenced NLR genes. First two components are shown and explain respectively 18 and 12% of the variance.

To confirm our calculations, we computed the correlations for π, π_s_ and *F*_ST_ at the CT loci between our data and a previous study, which used different plants from the same populations (Böndel *et al.*, 2015). There is a strong and significant correlation for π (R^2^ = 0.95, p = 3.7*10^−6^), π_S_ (R^2^ = 0.95, p = 5.8*10^−6^) and pairwise *F*_ST_ between populations (R^2^ = 0.94, p = 2.2*10^−16^; S Figure 1C-D). We could finally confirm the majority of SNPs in a subset of genes using Sanger sequencing (Supplemental Data 1), demonstrating the robustness of our SNP call approach and computation of diversity statistics for our pooled data.

### NLR genes show a wide range of diversity statistics

We find between 2,748 and 7,653 SNPs within each of the 14 sequenced populations. Across the set of 201 NLRs, 63.8 (± 0.48)% of SNPs are found on average to be non-synonymous, contrary to only 34 (± 3.26)% of non-synonymous SNPs at the CT genes. PCA analyses of the NLR SNPs show that most variation can be explained using the first two principal components, which reflect the geographical locations of the populations (Figure 1B). For each group, the median π is significantly higher for NLR than for CT genes (Figure 3A, p< 10^−5^). The reduced π values observed for the CT and NLR genes in southern mountain and coastal populations are indicative of the demographic consequences of the colonisation events that occurred during the species expansion southwards.

The π_N_/π_S_ values for most genes remain below one, indicative of purifying selection. However, NLR genes have significantly higher π_N_/π_S_ than CT genes (Figure 2B). Such higher π_N_/π_S_ values could indicate the occurrence of weak positive or balancing selection but also relaxed constraints at the NLRs. Mean and median π values are similar between CT loci and NLRs, but the variance is larger in NLRs. Six NLR genes show very large (median >0.02) π, and 15 genes show high π_N_/π_S_ (median > 1) (S Figure 2A).

**Figure 2.**
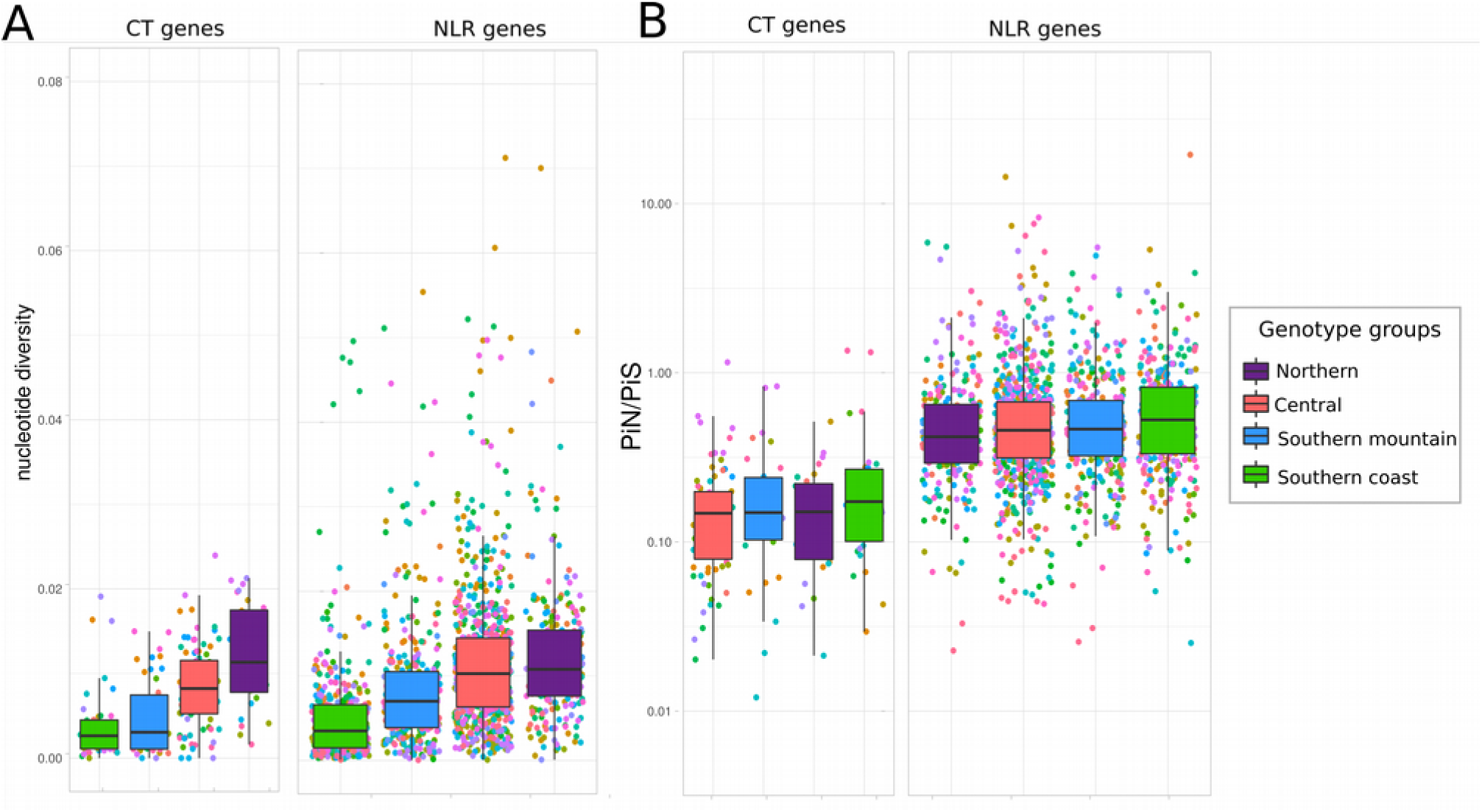
Population genetic statistics for NLR and CT loci A) Nucleotide diversity (π) for each gene, plotted per geographic group. B) Non-synonymous over synonymous nucleotide diversity (π_N_/π_S_) for each gene, plotted per group. Box plot colours match those of the geographic groups on Figure 1. Each dot represents a single gene, colours are assigned randomly.

**Figure 3.**
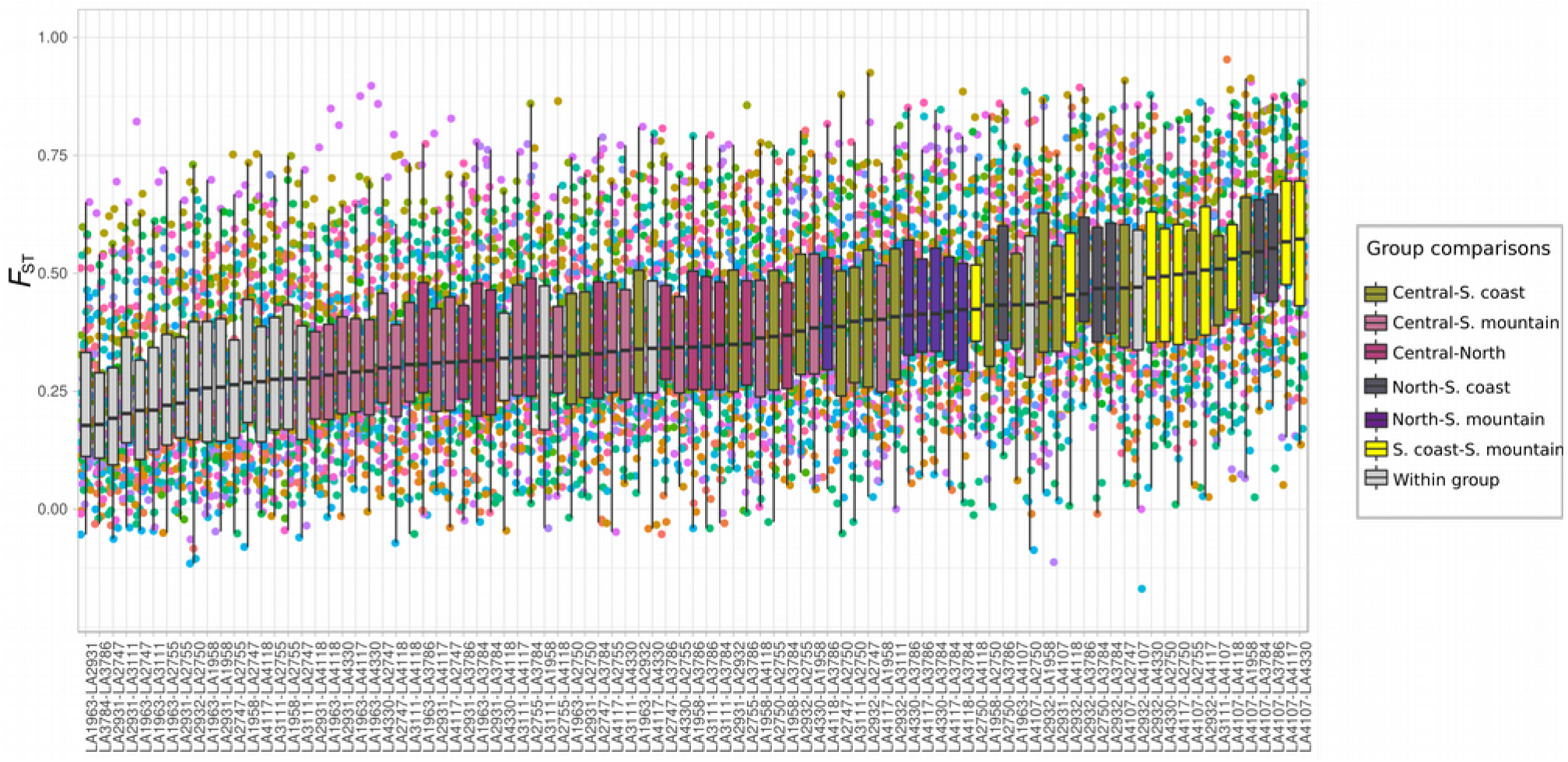
NLR Fixation index Fixation index (*F*_ST,_ y-axis), for each gene in each pairwise comparison between populations (x-axis). Colours of the boxes indicate the pairwise group comparisons. Each dot represents a single gene.

To compare the signatures of selection at the short time scale (polymorphisms within a species) with those at the longer time scale of divergence (between species), we compare, respectively, the π_N_/π_S_ within LA3111 to dN/dS calculated for our reference genome (LA3111) against *S. pennellii* LA0716. The dN/dS distribution over the NLRs does not differ from that at the CT loci (Böndel *et al.*, 2015) (t-test p = 0.17), nor when comparing between the different functional NLR clades (ANOVA, p = 0.6) (S Figure 3A). In the CT genes, which all have orthologs in *S. pennellii*, π_N_/π_S_ values are correlated to dN/dS (corr 0.65, p-value 0.02). This correlation is weaker at the NLRs for which orthologs in *S. pennellii* can be found (corr 0.33, p-value 0.004). Moreover, π_N_/π_S_ is significantly higher in NLRs which do not have any ortholog in *S. pennellii* than for the other NLRs (p-value = 0.003, S Figure 3B).

### Spatially heterogeneous selection pressure acting on different NLR functional classes

When NLRs are grouped by functional clades, we see that the CNL6 and NRC show very low π _N_/π_S_ and the newly identified clades (CNL20 and CNL21) show high values (S Figure 4A). Interestingly, contrasting patterns appear between the geographical groups (S Figure 4B). CNL11 shows the highest π_N_/π_S_ values in the coastal populations, whereas these values are lowest for CNL2 at the coast. Genes with π _N_/π_S_ > 1 differ between groups and populations, indicating that genes of the functional NLR clades are under different evolutionary pressures in the different geographical regions. NRCs appear quite conserved at both the phylogenetic time scale (between species) (low median dN/dS ratio for the LA3111 genome compared to *S. pennellii*, S Figure 3C) and at the polymorphism time scale (within species) (low median π_N_/π_S_, S Figure 4B). We calculated the fixation index (*F*_ST_) based on π for each gene between each pair of populations (Supplemental data 3). *F*_ST_ can be interpreted as a measure of genetic differentiation. We assure robustness of the calculations by using only 91 NLR genes with high and even coverage between all compared populations. Median *F*_ST_ values per NLR gene range between 0.12 and 0.7, with 17 genes having a median *F*_ST_ over 0.5 (S Figure 5). As expected, *F*_ST_ is lowest within geographic groups and highest between the coastal and the southern mountain populations (Figure 3).

**Figure 4.**
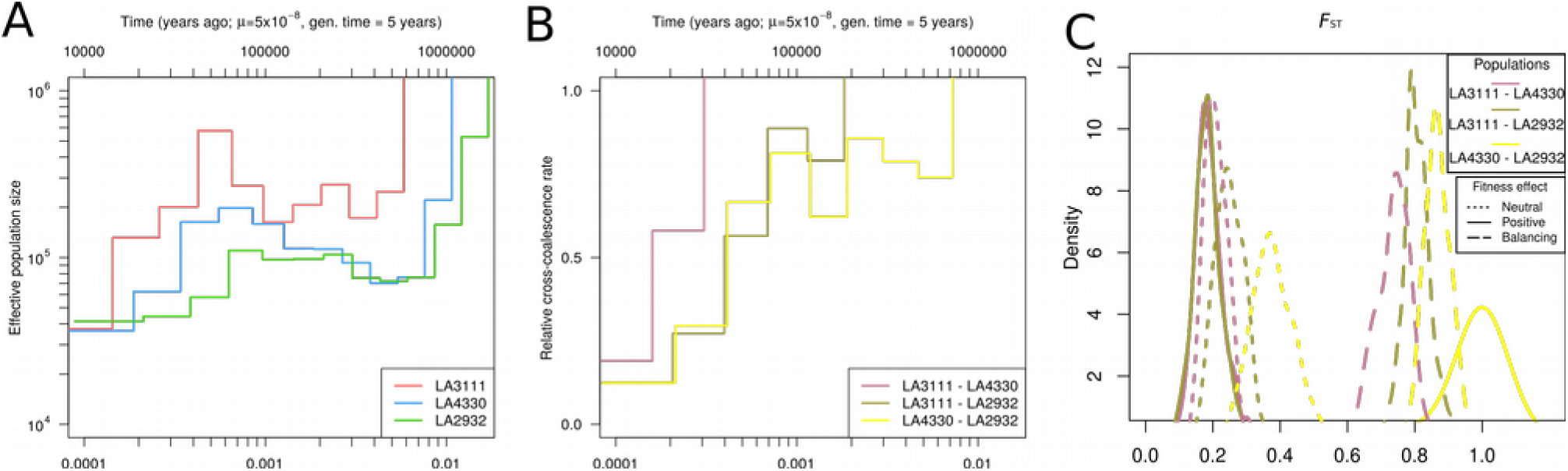
Historical demography reconstructions based on whole genome data of one individual from each of the central, southern coast and southern mountain populations. A) Effective population size (*N*_e_) through time estimations for central (LA3111; red line), southern coast (LA2932; green line) and southern mountain (LA4330; blue line) populations obtained with MSMC. Y-axis indicates the *N*_e_, x-axis the time in years ago (top) B) Estimation of the genetic divergence between pairs of populations through time: central-mountain (LA3111-LA4330; salmon line), central-coast (LA3111-LA2932; olive line) and mountain-coast (LA4330-LA2932; yellow line). The measures are based on the ratio between the cross-population and within-population coalescence rates (y-axis) as a function of time (x-axis). A rate of one indicates panmictic populations and rates of zero indicate fully separated populations. C) Genetic differentiation distributions (*F*_ST_) among the central, southern coast and southern mountain populations. Simulated genes evolve under neutrality in the central group and under either neutral, positive or balancing selection regimes in both southern populations following the colonization scenario and demography inferred with MSMC. *F*_ST_ (x-axis)is plotted against the observed density (y-axis), The comparisons are coloured as in B.

**Figure 5.**
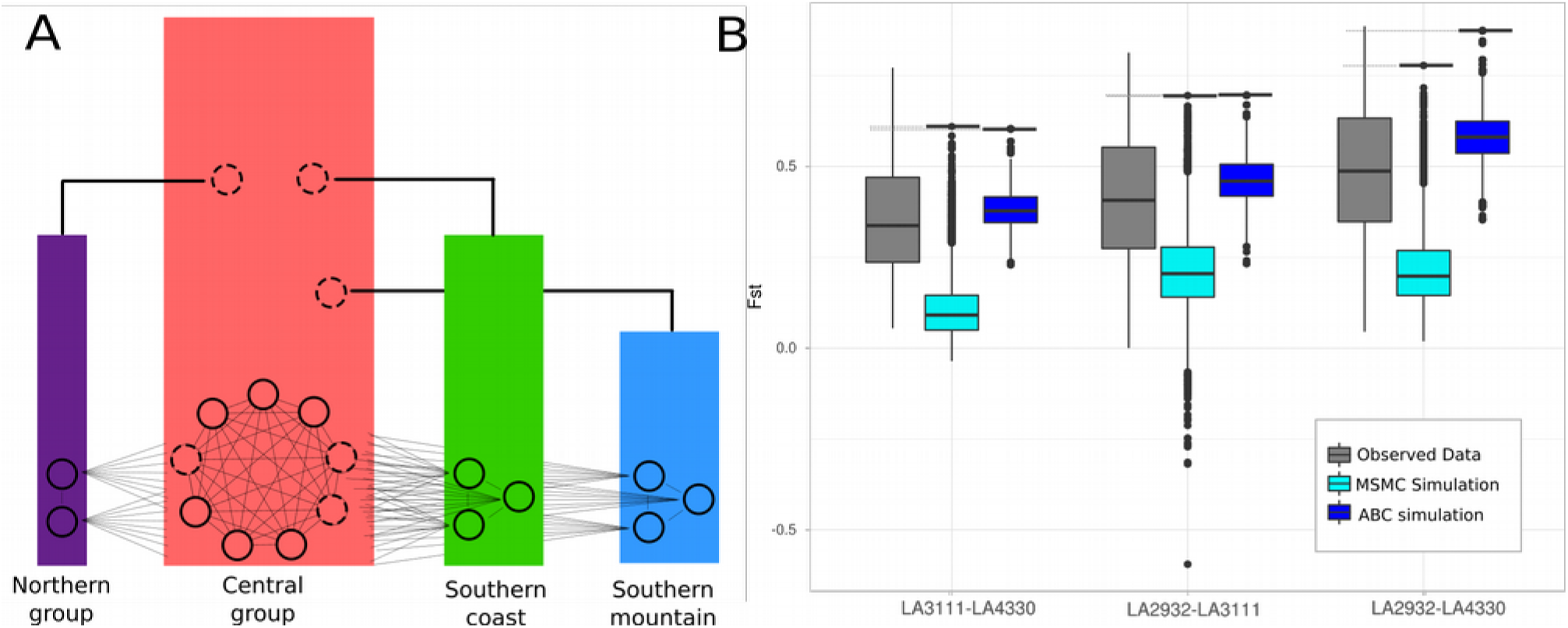
Coalescent simulations to identify *F*_ST_ cut-off values A) Coalescent model simulated for parameter estimation through the Approximate Bayesian Computation (ABC) approach. The model presents the same sampling for the empirical dataset with 14 populations from four regions (solid circles), as well as three unsampled “ghost” populations from the central region from which the populations of the other groups diverge. To illustrate that they also contribute to genetic diversity in the central population at present time they are presented twice in the figure (dashed circles). Populations evolve under the island model where migration among groups is smaller than migration within groups. B) Boxplots indicating the similarity between the observed data (grey) and our simulations based on ABC inference (dark blue) or the MSMC inference (turquoise). The black horizontal bars (and dotted extension) indicate the maximum simulated values. The simulations are based on 30,000 genes under the model inferred by MSMC or ABC. The maximum values obtained under the ABC model (top lines), are used as *F*_ST_ cut-offs for outlier selection.

### Genome-wide inference of the species’ past demographic history

We inferred the demographic history of three *S. chilense* populations LA3111 (central), LA2932 (southern coast) and LA4330 (southern mountain) using whole genome sequence data. We find consistent population expansion events for the three populations between 50 to 500 thousand years ago before reaching current *N*_e_, with a stronger expansion for the central group than for the other two populations (Figure 4A). Divergence estimations support that the central group is the area of origin of the species (Figure 4B). The species’ dispersal towards the new habitats occurred via two separate colonisation events around the Atacama desert: an older split between the central and coastal populations 0.2 to 1 million years ago, and a more recent divergence between the central and southern mountain populations, 30 to 150 thousand years ago. Note that the bottlenecks towards the south are relatively mild as *N*_e_ remains above 10^4^.

We tested the power to estimate known demographic histories with our genomic data. We simulated two (single-population) demographic scenarios for each population: one with a constant population size and one with a recent bottleneck event (S Figure 6B-C). Subsequently, we simulated two plausible scenarios derived from the interpretation of the observed data with both *N*_e_ changes and population splits including all three populations: one scenario limited to a single *N*_e_ change (i.e. bottleneck), and a more complex scenario with several *N*_e_ changes during the divergence processes (S Figure 6D-E). MSMC estimations from the simulated data verified the ability to recover known demographic parameters. We also use those simulations to compare the obtained demographic estimates from the empirical data. We find that the latter simulated scenario showed the best fitting to observed data (S Figure 6E).

**Figure 6.**
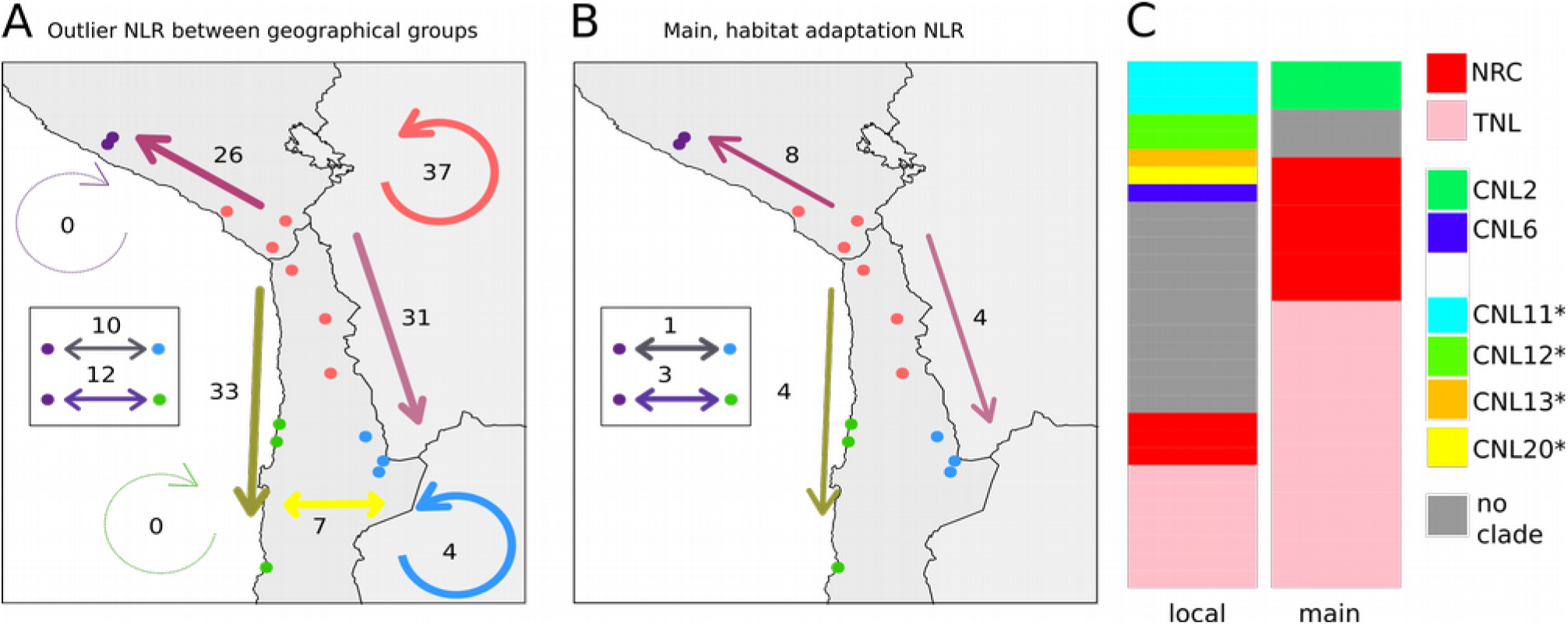
NLR genes under selection can be divided in main “habitat adaptation” and local “fine tuning” adaptation NLRs. A) Number of genes under selection (*F*_ST_ outliers) found between the four main geographical groups (straight arrows) as well as within each of the groups (circular arrows), when summing all individual outliers. NLRs under selection between the north and mountain or north and coast are indicated in the box. In total 53 genes can be identified, many are common to several geographic groups. B) Maps showing the number of main “habitat adaptation” NLRs between the different geographical groups. Habitat adaptation NLRs are defined as those that occur in more than one third of the possible population comparisons between the examined geographic groups. C) Functional clade assignment (as fraction) of the main adaptation and local fine tuning NLRs. Clades marked with an * are expected to be NRC-dependent

To confirm that under the inferred demography of *S. chilense, F*_ST_ statistics can be used as indicators of different selective pressures between populations we used forward simulations to generate polymorphism signatures of genes either under neutral, positive or balancing selection between populations during the population divergence with mild bottlenecks. Genes under positive or balancing selection in the southern populations can be differentiated from the neutral genes showing high value outliers in population pairwise *F*_ST_, in spite of the mild bottleneck effect that increases variance in *F*_ST_ distributions (Figure 4C). Studying low *F*_ST_ values for evidence of genes with similar selection pressures across populations is not powerful enough given our demographic history. We thus concentrate on high *F*_ST_ outliers between populations, which indicates novel and heterogeneous selective pressures (positive, balancing or relaxed constrains) in the derived populations (Charlesworth *et al.*, 1997).

### Defining *F*_ST_ cut-off values in a species-wide population structure

To study selection at NLRs over the whole species range (*e.g.* in our 14 populations which includes also a northern group of two populations) we additionally infer the past demographic history taking into account post-divergence migration by means of an ABC approach. We tested three models that include different scenarios for the divergence, while accounting for migration (Figure 5A). As observed summary statistics we use data at synonymous sites from all 91 NLRs and 14 CT loci to compute π per population and all pairwise *F*_ST_. In concordance with the results obtained with the whole-genome approach, the inference from ABC confirmed that the divergence of coast and mountain from the central group were two independent processes (Model 1; Supplementary Notes S4 - Figure1). This model showed strong support in five out of six rejection analyses performed (two rejection methods x three threshold values for simulations retained; Model 1 (Supplementary Notes S4 - Table1). As expected, posterior paramenter estimations showed higher *N*_e_ for populations from the central region. Lowest *N*_e_ value was estimated for south coastal populations. We also estimated higher gene-flow within the central group as well as among populations from the south mountain with the central group (Supplementary Notes S4, Table2).

We used the posterior distributions of the parameters based on the best supported model to simulate the pairwise *F*_ST_ between 14 populations for 30,000 loci (approximately the number of genes in the genome) with a mean gene size equal to that of our NLRs. Our inference yields a good fit to the observed values (S Figure 7). Especially when using the ABC, we were able to simulate median values that are very close to those of the observed data (Figure 5B). We can thus use the maximum of the simulated values as a conservative cut-off for *F*_ST_ based outlier detection.

### Revealing genes under selection as outlier loci: specific subgroups of NLRs evolve in each habitat

We identify the outlier NLRs as those whose *F*_ST_ values are found outside the simulated ranges for each pairwise population comparison (shown in Figure 6B for three population comparisons). We find a median of 7 NLRs to be outliers in all pairwise comparisons. In total 52 NLRs are found as outliers in at least one of the 91 possible pairwise comparisons (S Figure 8A). How often a gene is found as outlier in a pairwise comparison differs greatly. For example, eight NLRs appear as outliers in more than 15 pairwise comparisons, whereas six are identified only once or twice (S Figure 8B). When we sum the results per geographic group, we find a similar number of genes showing signatures of selection (due to genetic differentiation) in the southern coast or the southern mountain group and slightly less between the northern and the central group (Figure 6A). We also find NLRs under selection within the central group as well as some in the southern mountain group, but not within the northern or southern coastal groups.

### Main habitat adaptation NLRs and local adaptation NLRs belong to different functional classes

We now define NLRs that are under strong evolutionary pressure in multiple comparisons between the geographical groups as “main habitat-adaptation” genes. They are found to exhibit common selective pressure in several populations of the derived groups compared to the central group. We suggest that the pressure at these genes is shaped by global changes in habitat and/or pathogens during the early phase of colonisation. We analysed all pairwise *F*_*ST*_ comparisons between populations and find that 17 of the 52 outlier NLRs are main habitat-adaptation genes. These are outliers in more than 1/3 of the possible comparisons between two groups (Figure 6B). The remaining 35 genes that are under selection, only appear in few population comparisons between groups or only within a geographical group. We define these as “local adaptation” genes, presumably responsible for population level adaptation. These are clear outliers based on our neutral demographic model but do not exhibit habitat specific patterns, but rather exhibit an heterogeneous geographic mosaic of selection.

Looking at functional classes, main adaptation NLRs are more often TNLs or likely to belong to the NRC clade. The local adaptation genes are found to contain more individual NLRs that do not belong to a clade, belong to clades not part of the NRC-network, or are sensor NLRs (Figure 6C). By performing a randomisation procedure, we confirmed that the observed clade distribution of the NLRs under selection is unlikely to have arisen by chance. The observed number of major genes we find (17) is much larger than the expectation which has mean 3.5 (and C.I. [0.1-7.2]). Similarly, the observed fraction of NRC genes amongst the main habitat adaptation genes is five and larger than the expected one NRC (CI. [0.6-2.3]).

## Discussion

### NLR show sequence diversity within and between populations

NLR genes are important in plant defence responses and some have been shown to be under selection between different *Arabidopsis* species or populations (Mondragon-Palomino & Gaut, 2005; Bakker *et al.*, 2006). We used R-gene enrichment sequencing to investigate the extent of adaptation in the NLR family across wild populations of a non-model species, *Solanum chilense*.

We calculated synonymous and non-synonymous polyphormism statistics to asses possible selection on the NLRs. dN/dS ratios can be used to assess divergence of genes between species, and π_N_/π_S_ is the preferred statistic within species (Kryazhimskiy & Plotkin, 2008). High genetic diversity (observed as π and π_S_ values) is prevalent throughout the species. Between species diversity ratios (dN/dS) (*S. chilense – S. pennellii*) does not correlate with diversity ratios within *S. chilense* (π_N_/π_S_). This suggests recent positive selection is acting on the NLRs.

The π_N_/π_S_ ratio remains below 1 for the majority of the NLR genes in all populations, suggesting purifying selection and that the function of most NLRs is conserved within and between populations. Differences in diversity (and of the ratios) can be observed between previously defined genetic groups, with lower diversity in the derived groups. Yet, in all groups some NLRs exhibit high (non-synonymous) diversity, indicating that novel specitifties at NLRs appear and are picked up by natural selection as proposed in the NLR recylcing scenario (Holub, 2001). Note, *S. chilense* exhibits two previously undefined functional clades of NLRs in the *Solanaceae* (CNL 20 and CNL21), indicating the importance of birth and death process generating new NLRs with novel function at the phylogenic time scale (Michelmore & Meyers, 1998).

### Demographical inferences support two independent southward colinization processes in *S. chilense*

We implemented two demographic approaches that support a southward colonisation process already proposed by Böndel et al. (2015). This ocurred via two independent events over the last 200.000 years, one through the coast and the other across the highlands, resulting in two new sub-specific lineages in contrasting habitats. We find some discrepancies between ABC and MSMC in the divergence time estimations. These are expected given the differences in the approaches, model assumptions (*i.e.* considering or not migration) and the data used (*i.e.* set of genes vs. genome-wide) (Beichman *et al.*, 2017). In addition, gene exchange during divergence leads to an increment of variance of coalescence time among genes (Wakeley & Hey, 1997) causing discrepancies between population divergence and gene coalescence time estimations, especially for scenarios of small divergence times compared to *N*_e_ (Slatkin *et al.*, 2002).

Even taking into account intrinsic bias to the methods used, we consider that the two demographic approaches provide complementary evidences. We were able to generate a wide neutral distribution of genetic differentiation which is conservative enough to avoid false positives in the identification genes under selection. When using the ABC, we were able to simulate median values that are very close to those of the observed data. Furthermore, we provide evidence of the good fit of the demographic estimations to our data. For the MSMC method, we demonstrate using simulations that the high amount of nucleotide diversity and recombination rate found in *S. chilense* (Roselius *et al.*, 2005).

### NLR change selection within and between habitats

We used demographic inference to establish a neutral distribution of the genetic differentiation to define outlier NLRs that change selective pressure between populations. We found that during the intra-specific differentiation NLRs not only change selection between different geographical groups, but also regularly between populations within the same region, especially in the Central group. Böndel et al. (2015) already found the central group to be genetically more diverse and noted that it should maybe not be treated it as a single panmictic unit because its relative high climatic heterogenity. We postulated that the coastal and southern environments differ in their biotic factors from the central region (Stam *et al.*, 2017). In the Coastal region, we expected to observe a lack of selection on NLRs as we assumed the arid environment would be void of phytopathogens. Contrary to our hypothesis, our data show selection towards the coast and thus indicate that pathogens are historically present. This could for example be due to seasonally running rivers as well as a regularly occurring sea-fog phenomenon in the early morning (Cereceda & Schemenauer, 1991).

### Major and local adaptation NLR

Our results allowed us to separate major habitat adaptation NLRs from local adaptation NLRs. The 17 major habitat adaptation show changes of selection throughout the species’ distribution, with different major genes between each geographical group. Habitat adaptation NLRs more often belong to the class of helper-NLRs, called NRC (Wu *et al.*, 2017a), as well as to the TNL. NRCs are hypothesised to be under strong purifying selection due to their central role (hub) in the NLR-signaling network (Wu *et al.*, 2018). Indeed, NRCs showed low dN/dS values and overall, π_N_/π_S_ is low in the NRCs. High fixation (*F*_*ST*_) for between populations for some NRC, indicates that minor changes in individual hub proteins could also be under strong selection. In *A. thaliana* RPW8-like NLRs, ARD1 and NRG1, function as helper NLRs for TNLs and are required for functioning of NLRs against several well studied pathogens (Brendolise *et al.*, 2018; Qi *et al.*, 2018; Castel *et al.*, 2019). In our study, RPW8-like genes are not detected as outliers. This could be explained by the fact that ARD1 and NRG1 have no clear homologues in the *Solanum* genus and thus that TNL signaling in this genus is likely to function differently, possibly with a subclade of TNLs taking over the function of hubs.

Local adaptation NLRs are more often not assigned to known functional clusters, or smaller clades, like the newly defined CNL20, suggesting that new clades are involved in local fine tuning of the defence responses, generating a geographic mosaic (Thompson, 2005) of NLR variants that have co-evolved with local pathogens. It is know that the NRC-dependent R-gene Pto (and other genes of the Pto signalling network), indeed shows such large allelic variation and is under balancing selection within different wild tomato species, including *S. chilense* (Rose *et al.*, 2007, 2011).

### Scenarios leading to two-tiered selection of NLR in new habitats

Changes in the NRC-clade dependent defence response thus rely on co-evolution of both the sensor and the helper NLR, rather than the evolution of the sensor alone. We hypothesize that within *S. chilense* each NRC co-evolves as a helper NLR with a specific set of sensor NLRs. In experimental evolution in yeast, major evolutionary and functional novelty has been shown to occur by changes in the hubs of a gene network (Koubkova-Yu *et al.*, 2018). The main genes underlying habitat adaptation are often “helpers” and do not on their own provide a specific recognition of the newly encountered pathogens (new species or genera), but improve signalling processes. Several single non-synonymous mutations have been shown to result in gain of function of NRC1 for downstream signalling activity (Sueldo *et al.*, 2015). Moreover, NLR functioning is known to be dependent on temperature (Cheng *et al.*, 2013) and other abiotic stresses (Ariga *et al.*, 2017). In *S. chilense* different NRCs could, for example be responding to different temperatures between the coast and the mountains.

In a fixed habitat, genes that are well connected in the defence gene network would be expected to be under strong functional constraints (purifying selection). Such selection has, for example been described for the NRC-independent I2 gene in *S. pimpinellifolium* (Couch *et al.*, 2006). In newly colonised habitats, selection at these genes could be resulting from two possible scenarios. 1) The new mutations at the main habitat-adaptation genes enable their binding with different and previously unbound sensors or new binding abilities under different abiotic conditions. This scenario would explain the occurrence of new positive or balancing selection at the helper genes in the derived habitats, and that different NRC genes are under selection in the three new habitats. 2) The helper genes are under relaxed constraint because the associated sensor genes are not necessary as their specific associated pathogens are absent in the new habitat. The sensor could become non-functional, so that helper genes are free to evolve neutrally or even develop novel beneficial functions (neo-functionalisation), that are selected for in subseqtent generations. In both scenarios sensors NLR can freely evolve to optimizing the detection of the newly encountered pathogens in specific localities and would coevolve rapidly with the pathogens. Together, this would lead to the observed two-tiered selection process.

## Conclusions

Our work represents a first step in studying the dynamics of NLR evolution across space and across the gene/plant defence network at the population level. Our results strengthen the view that NLRs do not evolve on their own to sense/recognise pathogen molecules, but their evolution is constrained by their interaction with other genes in the network. Future work on reliable identification of functional R genes, as well as the effectors of natural pathogens present in the different populations, will allow us to study the population genetics of direct effector-target interactions (Terauchi & Yoshida, 2010) and thus provide insight into the molecular factors shaping the different plant-pathogen coevolutionary dynamics in nature.

## Supporting information

Supplemental Item 1

Supplemental Item 2

Supplemental Item 3

## Acknowledgements

RS was supported by the Alexander von Humboldt foundation. Genome resequencing was funded by DFG grant TE 809/7-1 to AT. GSA acknowledges the TUM University Foundation Fellowship. We thank Christine Wurmser (NGS@TUM) for help with the re-sequencing and Christopher Huptas and Mareike Wenning (NGS@TUM) for help with the NLR sequencing. We also thank the TGRC at UC Davis (USA) for plant material, and Anja Hörger, Tetyana Nosenko and Wolfgang Stephan for feedback on the manuscript.

## Notes

#### Summary of Updates

We made several major improvements: - We improve our demographic modeling of Solanum chilense. We use Approximate Bayesion Computation to calculate the expected Fst range for each pairwise population comparison to estimate a more reliable and pair-specific cut-off. - We use forward simulations to prove that under our demographic scenario change to Fst is a reliable indicator for changes in selective pressure from neutral evolution in the central region to positive or balancing selection in the newly colonized habitat. - We define major habitat adaptation genes and local fine tuning genes, based on how often genes change evolutionary pressure between populations. Genes that change pressure between many populations, e.g. throughout the whole species range are major habitat adaptation genes. Interestingly, we found those to belong more often than expected to the NRC clade of NLRs, thus genes central in the NLR network also change evolutionary pressure

