## Supplemental Item 1 for "Subsets of NLR genes drive adaptation of tomato to pathogens during colonisation of new habitats": Sanger.pdf

### Sequencing results / quality control

CT166 in LA3111, 1/1 TP, 0 FP

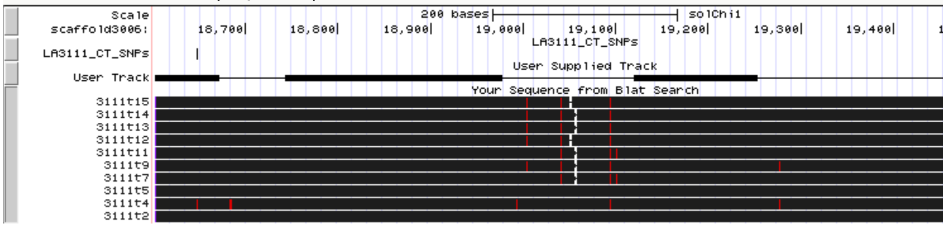


CT192 in LA2755, 18/18 TP, 2 FP

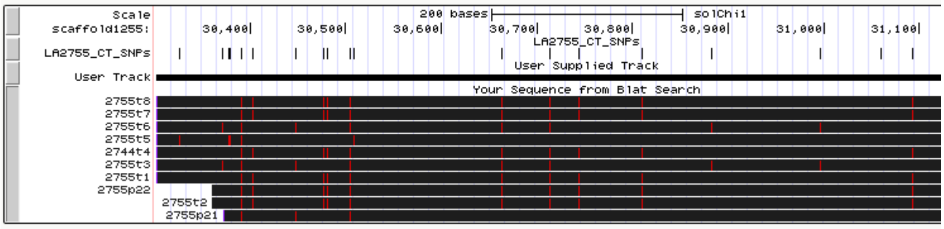


SOLCI000037500 in LA2931, 20/21 TP, 1 FN, 5 FP,

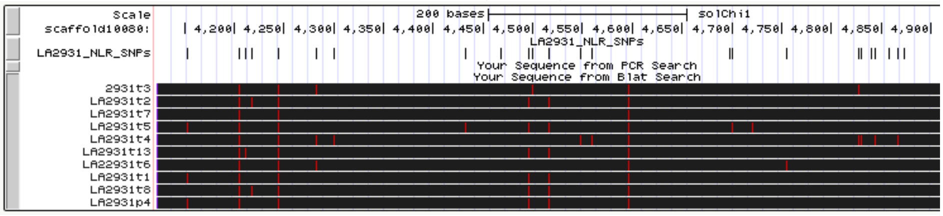


SOLCI002113000 in LA3111, 23/25 TP, 2 FN, 1 FP

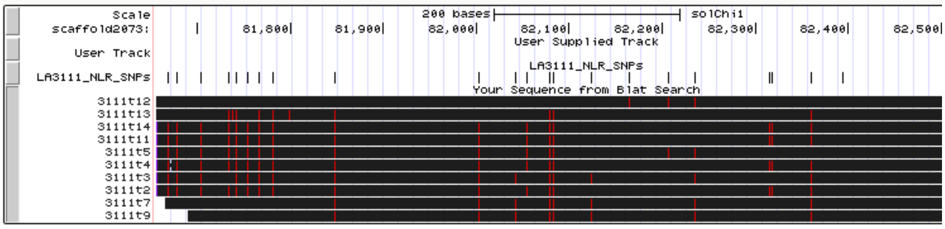


S Data, Sanger Sequencing

Genome Browser Screenshots showing from top to bottom, two CT and two NLR gene fragments. Each panel shows in the top half the genomic position on the scaffolds the identified SNPs in the respective population (as vertical black dashes) and in the lower half black bars indicating the Sanger sequenced region, with the actual found SNPs relative to the reference as red vertical dashes. Each black bar represents an individual plant. TP: True Positive, FN; False Negative, FP: False Positive None of the cloned fragments of CT166 have any SNPs, indicating absence of false negatives. CT192 has a high number of SNPs (18) at the 3' end in LA2755 of which we recover 100%, including two singletons. We also cloned parts of two NLR genes, SOLCI000037500 from LA2931 and SOLCI002113000 from LA3111. We find 21 and 25 SNPs, respectively, though 1 or 2 false negatives occurred
