## Supplementary Figures for "Subsets of NLR genes drive adaptation of tomato to pathogens during colonisation of new habitats"

### Supplemental Figures

S Figure 1

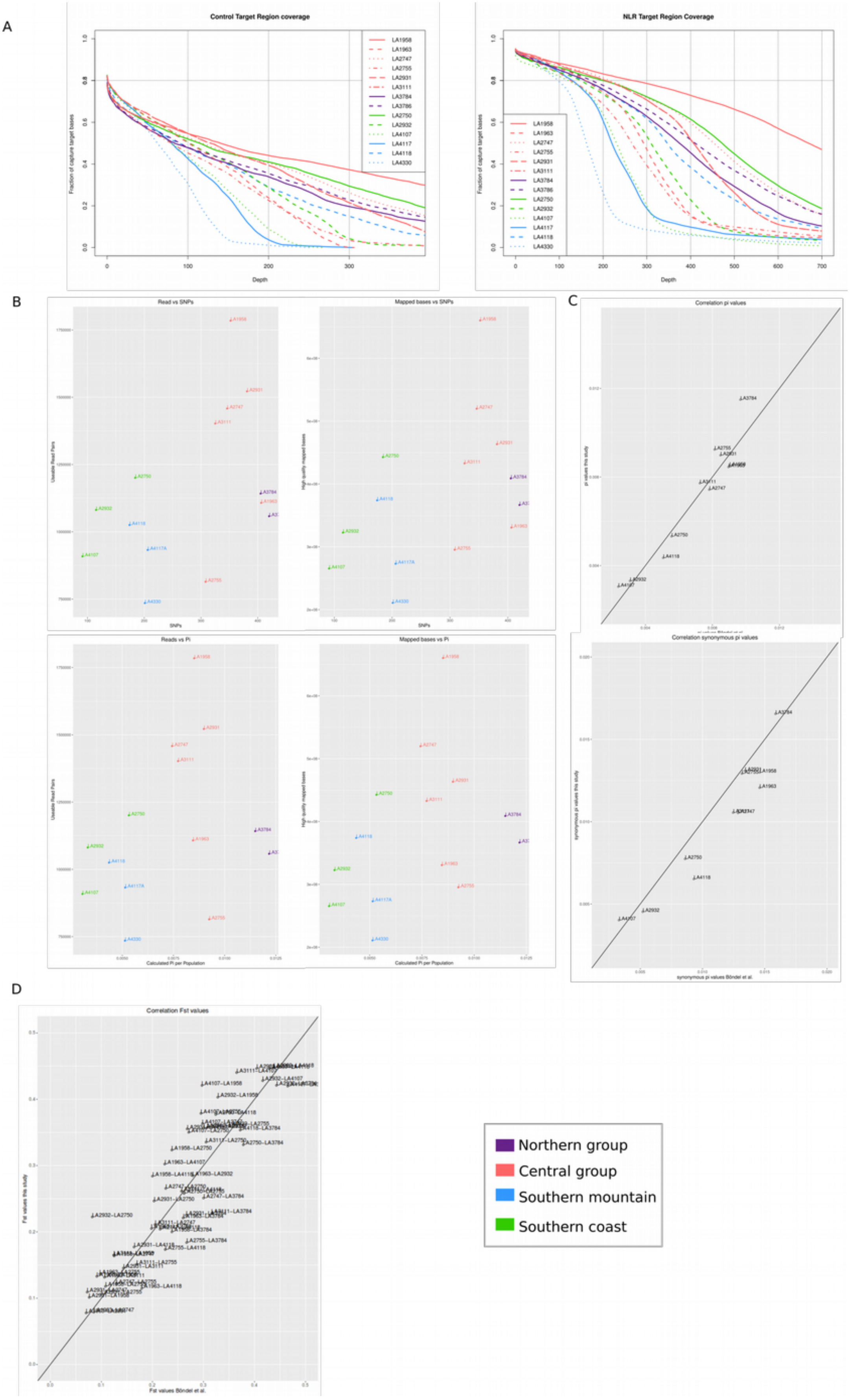

S Figure 1

A) Coverage plots for enrichment sequencing. The x-axis shows the depth of coverage, ranging from 0 to 400 or 700 for CT or NLR genes respectively. The y-axis shows the fraction of the targeted region (*e.g.* the part of the genome that was targeted by the probes) that has the indicated coverage or higher.

B) Number of sequenced reads (left) or high quality mappable read pairs (right) plotted against the total number of SNPs per sample (top) or π per population.

C) Calculated values for π (top) and $\pi_S$ (bottom) from this study (y-axis), plotted against the values obtained by Böndel et al. (x-axis). The studies used the same geographical populations but different individual plants.

D) Calculated values for $F_{ST}$ in pairwise population comparisons in this study (y-axis) plotted against the values obtained by Böndel et al. (x-axis). The studies used the same geographical populations but different individual plants.

S Figure 2

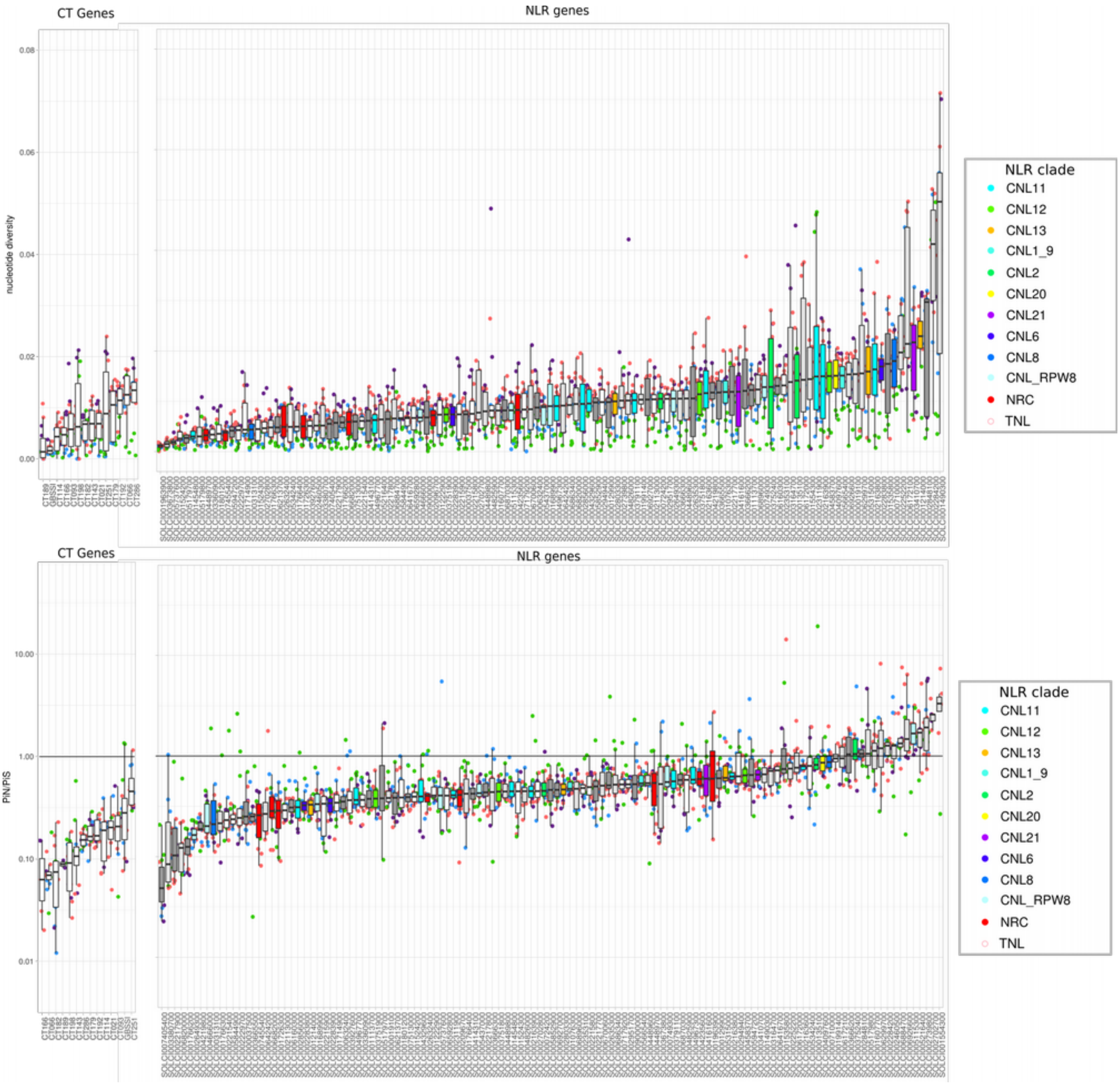


S Figure 2

A) Per gene overview for nucleotide diversity (π, top) and the non-synonymous over synonymous ratio ($\pi_N/\pi_S$, bottom). The distribution of values for NLRs (right) does in the majority of cases not exceed the range observed in CT genes (left). However, several genes with higher values can be observed, as well as individual outliers of a gene in one or several populations. Each point represents a single gene in one population, coloured by geographic group. The boxplots show median and upper and lower quartiles and the NLRs are coloured per cluster.

S Figure 3

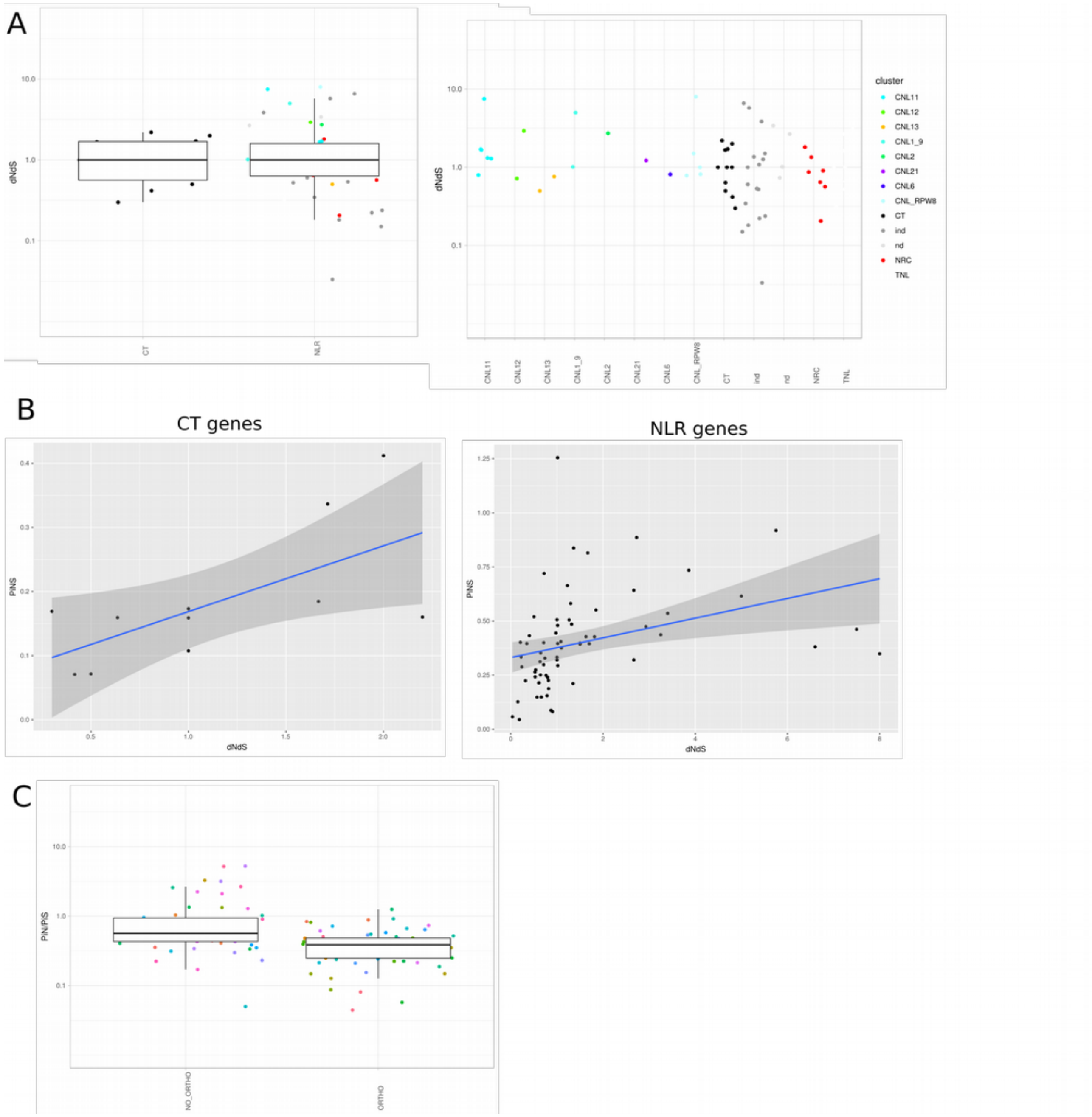


S Figure 3

A) dN/dS ratios for 14 CT and 91 NLR genes with orthologs in *S. pennellii*. The dN/dS is computed from the reference genome of LA3111. Data are split in CT and NLR (left panel) or plotted for each gene class individually (right panel). Each dot represents one gene and the dots are color coded after the functional clade (or cluster) to which they below. The dN/dS distribution over NLRs does not differ from that at CT loci (t.test p = 0.17) nor when comparing between the different functional clades (ANOVA, p = 0.6).

B) Correlations between $\pi_N/\pi_S$ for population LA3111 and dN/dS (from the reference genome LA3111) for CT and NLR genes (CT genes correlation corr 0.65, p-value 0.02, for NLRs corr 0.33, p-value 0.004).

C) Difference in $\pi_N/\pi_S$ (for population LA3111) between NLRs which do not have any ortholog in *S. pennellii* and those that do (right), t.test p-value = 0.003.

S Figure 4

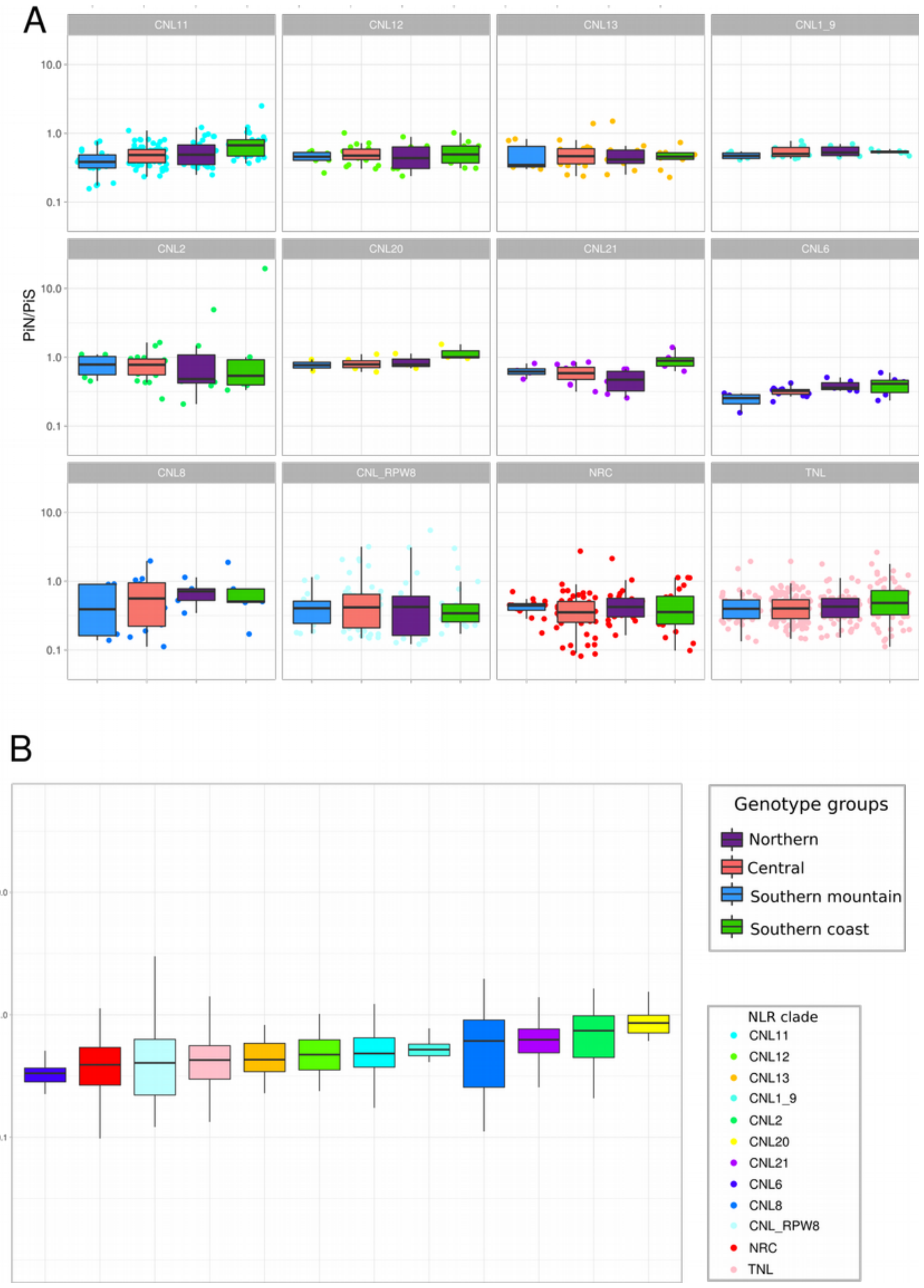


S Figure 4

A) $\pi_N/\pi_S$ for NLR classes over the different geographic group show variable patterns. Significant differences can be observed in clades CNL11, CNL13, CNL2, CNL21 and CNL6 (ANOVA, $p < 10^{-5}$ after posthoc testing).

B) Different NLR functional classes show large variation in $\pi_N/\pi_S$ over the whole species.

S Figure 5

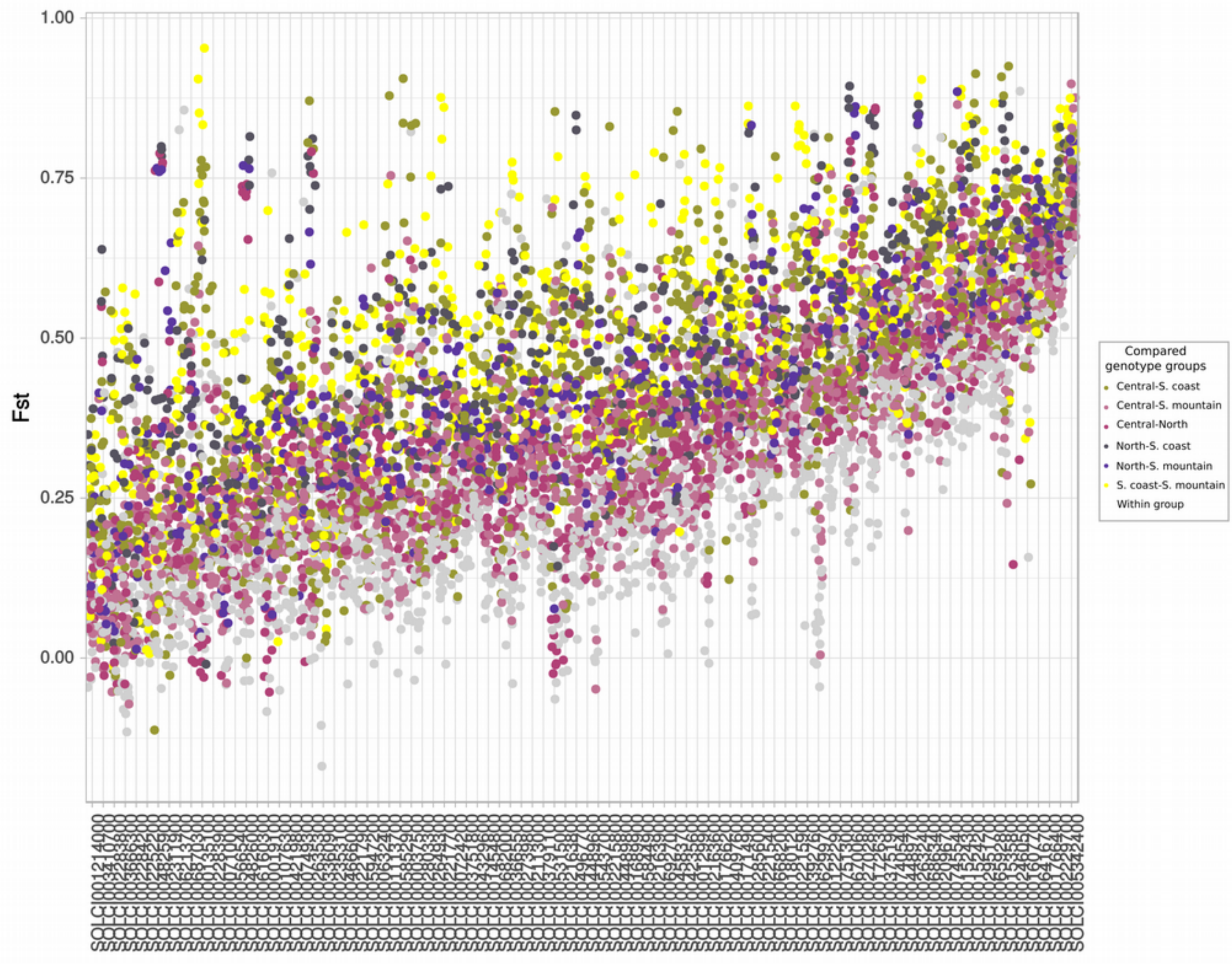


S Figure 5

C) Pairwise population $F_{ST}$ values (y-axis) are plotted for each NLR gene (x-axis) for each of the 91 possible comparisons. Each comparison is coloured based on the group  of origin of each populationinvolved..

S Figure 6

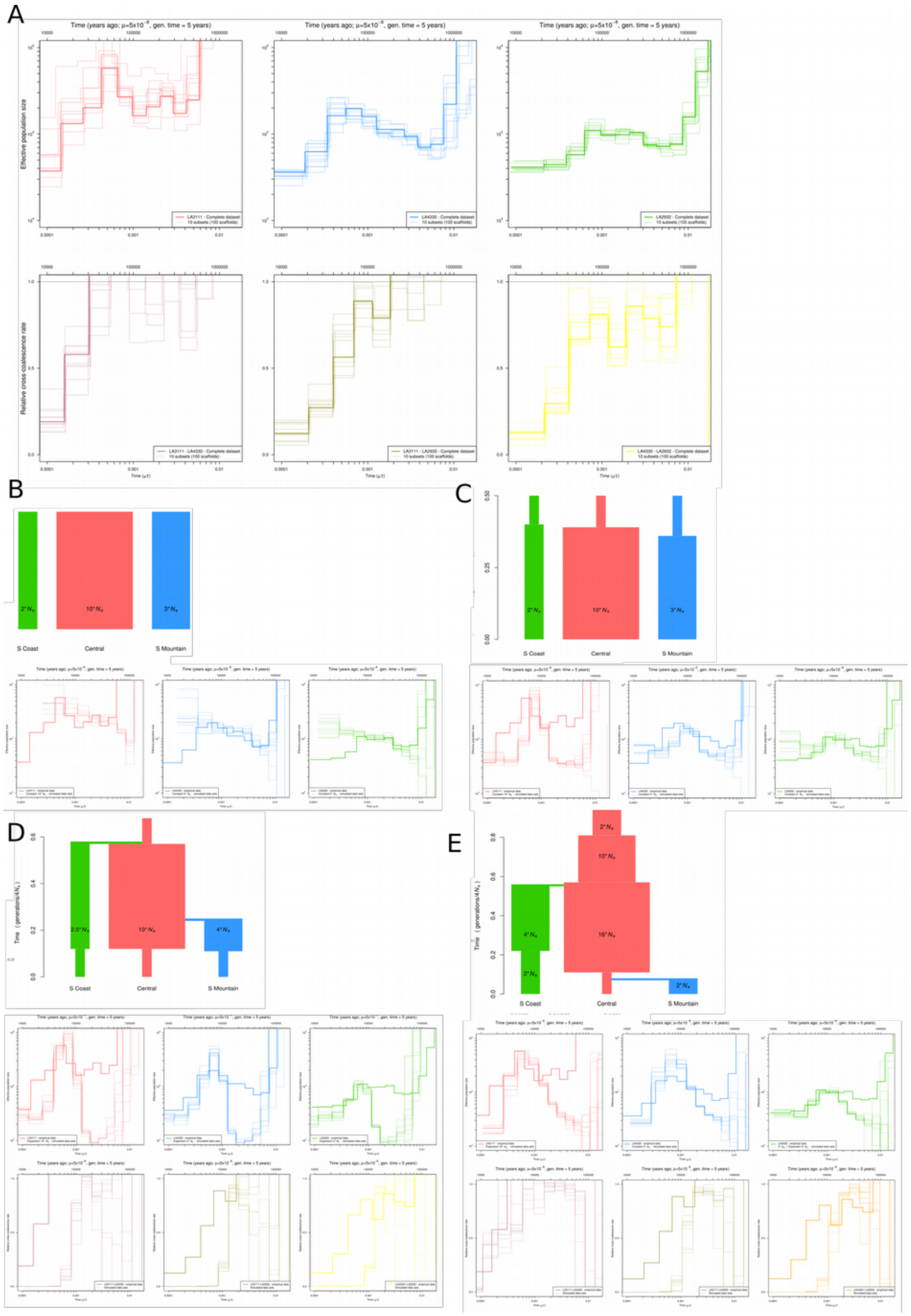

A
Time (years ago; μ=5x10⁻⁸, gen. time = 5 years)
Effective population size
Relative cross-coalescence rate
LA3111 - Complete dataset
LA4330 - Complete dataset
LA2932 - Complete dataset
10 subsets (100 scaffolds)
LA3111 - LA4330 - Complete dataset
LA3111 - LA2932 - Complete dataset
LA4330 - LA2932 - Complete dataset
Time (μ t)
B
2*Ne
10*Ne
3*Ne
S Coast
Central
S Mountain
C
2*Ne
10*Ne
3*Ne
S Coast
Central
S Mountain
D
Time (generations/4Ne)
2.5*Ne
10*Ne
4*Ne
S Coast
Central
S Mountain
E
Time (generations/4Ne)
2*Ne
10*Ne
4*Ne
16*Ne
2*Ne
2*Ne
S Coast
Central
S Mountain

S Figure 6

A) Confidence intervals of the demographic estimations obtained with MSMC. Upper row corresponds to the effective population size estimations of central (LA3111; red line), southern coastal (LA2932; green line) and southern mountain (LA4330; blue line) populations. Bottom row corresponds to the estimations of the genetic divergence between pairs of populations through time. Central-mountain (LA3111-LA4330; salmon line), central-coast (LA3111-LA2932; olive line) and mountain-coast (LA4330-LA2932; yellow line).

The solid lines represent the estimations obtained with the complete dataset (200 scaffolds of at least 200kb each), whereas the dashed lines represent estimations based on 10-times repeated random sub-sampling of 100 scaffolds (each at least 200kb).. Genetic divergence between populations is measured as relative cross-coalescence rates (y axis) as a function of time (x axis) with 1 indicating well mixed populations and 0 indicating fully separated populations.

B) Effective population size ($N_e$) estimations for individual simulations of constant $N_e$ scenarios.

The simulations were based on the current $N_e$ value estimated with MSMC for the empirical genome data. The solid line represents the complete dataset, whereas the dashed lines represent the estimations for 10 independent simulations. Times on the lower x-axes are given in units of divergence per base pair and times on the upper x-axes are given in years before present.

C) Effective population size ($N_e$) estimations for individual simulations of past expansion of $N_e$ scenarios.

The simulations were based on the current $N_e$ and the time of population expansion values estimated with MSMC for the empirical genome data. The solid line represents the complete dataset, whereas the dashed lines represent the estimations for 10 independent simulations. Times on the lower x-axes are given in units of divergence per base pair and times on the upper x-axes are given in years before present.

D) and E) Effective population size ($N_e$) and genetic divergence estimations for individual simulations of a demographic scenarios inferred from the MSMC results.

The simulations were based on the current $N_e$, the time of population expansion, and split times values estimated with MSMC for the empirical genome data. The solid line represents the complete dataset, whereas the dashed lines represent the estimations for 10 independent simulations. The upper row corresponds to the effective population size estimations for central (LA3111; red line), coastal (LA2932; green line) and mountain (LA4330; blue line) populations. The bottom row corresponds to the estimations of the genetic divergence between pairs of populations through time. Central-mountain (LA3111-LA4330; salmon line), central-coast (LA3111-LA2932; olive line) and mountain- coast (LA4330-LA2932; yellow line). Times on the lower x-axes are given in units of divergence per base pair and times on the upper x-axes are given in years before present.

S Figure 7

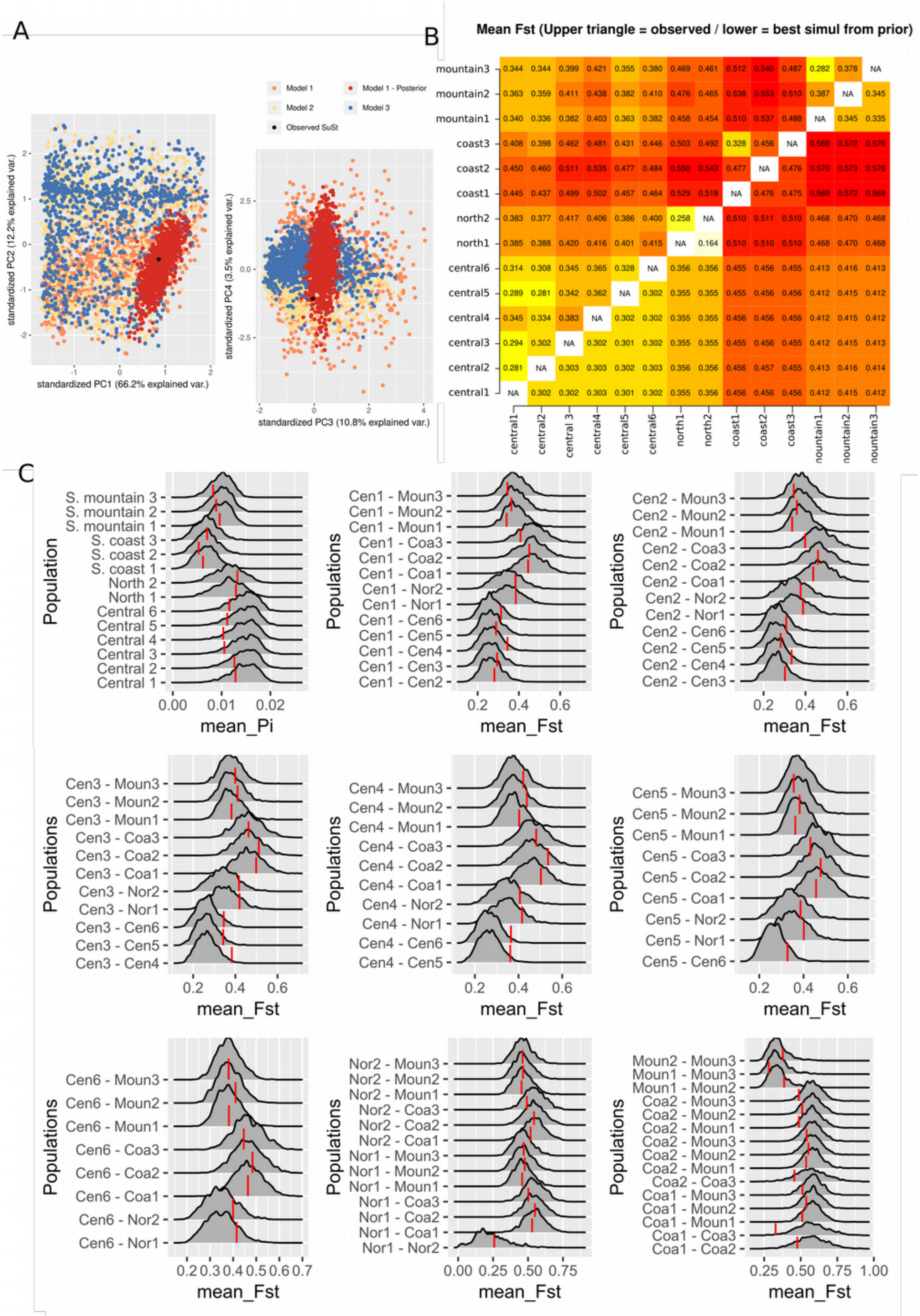
A
Model 1
Model 1 - Posterior
Model 2
Model 3
Observed SuSt
standardized PC2 (12.2% explained var.)
standardized PC1 (66.2% explained var.)
standardized PC4 (3.5% explained var.)
standardized PC3 (10.8% explained var.)
B
Mean Fst (Upper triangle = observed / lower = best simul from prior)
mountain3 0.344 0.344 0.399 0.421 0.355 0.380 0.469 0.461 0.512 0.540 0.487 0.282 0.378 NA
mountain2 0.363 0.359 0.411 0.438 0.382 0.410 0.476 0.465 0.538 0.553 0.510 0.387 NA 0.345
mountain1 0.340 0.336 0.382 0.403 0.363 0.382 0.458 0.454 0.510 0.537 0.488 NA 0.345 0.335
coast3 0.408 0.398 0.462 0.481 0.431 0.446 0.503 0.492 0.328 0.456 NA 0.569 0.572 0.570
coast2 0.450 0.460 0.511 0.535 0.477 0.484 0.550 0.543 0.477 NA 0.476 0.570 0.573 0.570
coast1 0.445 0.437 0.499 0.502 0.457 0.464 0.529 0.518 NA 0.476 0.475 0.569 0.572 0.569
north2 0.383 0.377 0.417 0.406 0.386 0.400 0.258 NA 0.510 0.511 0.510 0.468 0.470 0.468
north1 0.385 0.388 0.420 0.416 0.401 0.415 NA 0.164 0.510 0.510 0.510 0.468 0.470 0.468
central6 0.314 0.308 0.345 0.365 0.328 NA 0.356 0.356 0.455 0.456 0.455 0.413 0.416 0.413
central5 0.289 0.281 0.342 0.362 NA 0.302 0.355 0.355 0.455 0.456 0.456 0.412 0.415 0.412
central4 0.345 0.334 0.383 NA 0.302 0.302 0.355 0.355 0.456 0.456 0.456 0.412 0.415 0.412
central3 0.294 0.302 NA 0.302 0.301 0.302 0.355 0.355 0.455 0.456 0.455 0.412 0.415 0.413
central2 0.281 NA 0.303 0.303 0.302 0.303 0.356 0.356 0.456 0.457 0.455 0.413 0.416 0.414
central1 NA 0.302 0.302 0.303 0.301 0.302 0.355 0.356 0.456 0.456 0.456 0.412 0.415 0.412
central1 central2 central 3 central4 central5 central6 north1 north2 coast1 coast2 coast3 nountain1 nountain2 nountain3
C
Population
S. mountain 3
S. mountain 2
S. mountain 1
S. coast 3
S. coast 2
S. coast 1
North 2
North 1
Central 6
Central 5
Central 4
Central 3
Central 2
Central 1
mean_Pi
Populations
mean_Fst
Cen1 - Moun3
Cen1 - Moun2
Cen1 - Moun1
Cen1 - Coa3
Cen1 - Coa2
Cen1 - Coa1
Cen1 - Nor2
Cen1 - Nor1
Cen1 - Cen6
Cen1 - Cen5
Cen1 - Cen4
Cen1 - Cen3
Cen1 - Cen2
Cen2 - Moun3
Cen2 - Moun2
Cen2 - Moun1
Cen2 - Coa3
Cen2 - Coa2
Cen2 - Coa1
Cen2 - Nor2
Cen2 - Nor1
Cen2 - Cen6
Cen2 - Cen5
Cen2 - Cen4
Cen2 - Cen3
Cen3 - Moun3
Cen3 - Moun2
Cen3 - Moun1
Cen3 - Coa3
Cen3 - Coa2
Cen3 - Coa1
Cen3 - Nor2
Cen3 - Nor1
Cen3 - Cen6
Cen3 - Cen5
Cen3 - Cen4
Cen4 - Moun3
Cen4 - Moun2
Cen4 - Moun1
Cen4 - Coa3
Cen4 - Coa2
Cen4 - Coa1
Cen4 - Nor2
Cen4 - Nor1
Cen4 - Cen6
Cen4 - Cen5
Cen5 - Moun3
Cen5 - Moun2
Cen5 - Moun1
Cen5 - Coa3
Cen5 - Coa2
Cen5 - Coa1
Cen5 - Nor2
Cen5 - Nor1
Cen5 - Cen6
Cen6 - Moun3
Cen6 - Moun2
Cen6 - Moun1
Cen6 - Coa3
Cen6 - Coa2
Cen6 - Coa1
Cen6 - Nor2
Cen6 - Nor1
Nor2 - Moun3
Nor2 - Moun2
Nor2 - Moun1
Nor2 - Coa3
Nor2 - Coa2
Nor2 - Coa1
Nor1 - Moun3
Nor1 - Moun2
Nor1 - Moun1
Nor1 - Coa3
Nor1 - Coa2
Nor1 - Coa1
Nor1 - Nor2
Moun2 - Moun3
Moun1 - Moun3
Moun1 - Moun2
Coa2 - Moun3
Coa2 - Moun2
Coa2 - Moun1
Coa2 - Moun3
Coa2 - Moun2
Coa2 - Moun1
Coa2 - Coa3
Coa1 - Moun3
Coa1 - Moun2
Coa1 - Moun1
Coa1 - Coa3
Coa1 - Coa2

S Figure 7

Assessment of the accuracy of the demographic inference using ABC. Verification of the simulated summary statistics (π and $F_{ST}$ at synonymous sites) from the prior and posterior parameter distribution under an island model with isolation-migration scenario with 14 populations.

A) PCA of the summary statistics.

B) Matrix showing the observed mean $F_{ST}$ values from our NLR dataset (upper triangle) and the simulated values derived from our ABC-method (lower triangle).

C) Density plots obtained with 30,000 simulations of the summary statistics under the posterior parameter estimation. Vertical red lines indicate the observed values.

S Figure 8

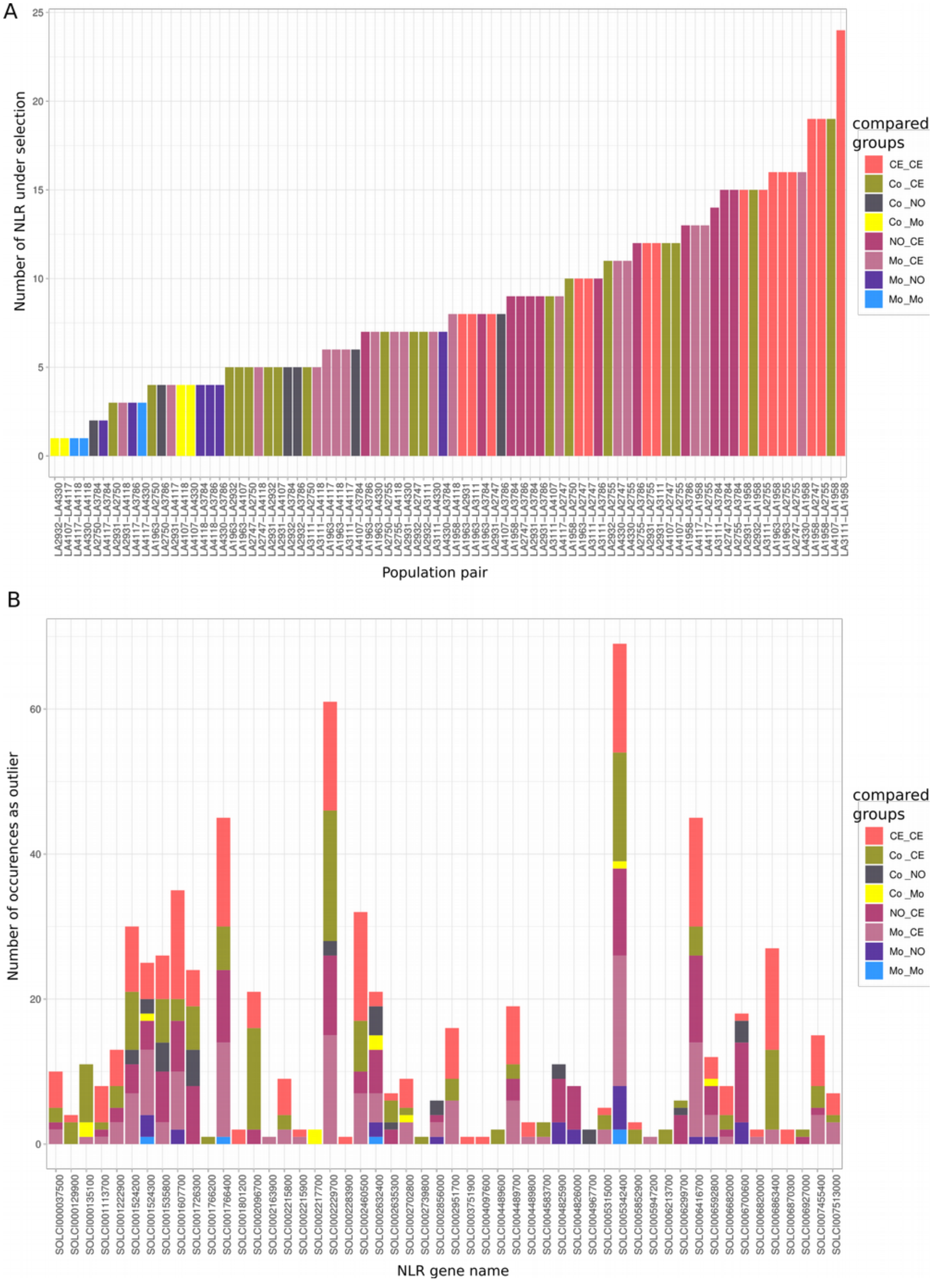
A
Number of NLR under selection
Population pair
compared groups
CE_CE
Co_CE
Co_NO
Co_Mo
NO_CE
Mo_CE
Mo_NO
Mo_Mo
B
Number of occurences as outlier
NLR gene name
compared groups
CE_CE
Co_CE
Co_NO
Co_Mo
NO_CE
Mo_CE
Mo_NO
Mo_Mo

S Figure 8

The number of NLR that are classified as an outlier based on the maximum simulated $F_{ST}$ value for each of the possible pairwise population comparisons. The bars are coloured based on the geographic groups to which the populations belong.

B) Count data (y axis) showing how often a certain NLR (x axis) is categorised as an outlier based on its $F_{ST}$ value. Counts are summed for each pair-wise comparison between populations.

(CE =central, NO = north, Mo = southern mountain, Co = southern coast).

Supplemental Data Item 1: Mapping QC stats

Supplemental Data Item 2: Summary statistics, NLR population genetics

Supplemental Data Item 3: Fst statistics per gene
