## Supplemental Data 1 for "Subsets of NLR genes drive adaptation of tomato to pathogens during colonisation of new habitats"

**STAM ET AL NLR EVOLUTION SUPPLEMENTARY NOTES**

**S1: PLANT & DNA MATERIAL**

We ordered seeds from 14 populations (e.g. a collection of interbreeding specimens collected at one geographical location, stored under one accession number) of *S. chilense* from the TGRC (UC Davis, California, USA, http://tgrc.ucdavis.edu) and grew 10 plants for each population in our glasshouse (20°C, 16h light). As shown in previous work on this species, the seed set per accession represents a good sample of the original genetic diversity in a population (Arunyawat *et al.*, 2007; Tellier *et al.*, 2011a; Böndel *et al.*, 2015). The chosen populations had accession numbers: LA3111, LA4330, LA2932, LA1958, LA1963, LA2747, LA2755, LA2931, LA3784, LA3786, LA2750, LA4107, LA4117(A), LA4118. These accessions were collected in different years and were multiplied different number of times at TGRC to produce seeds by random crossing. However, it is shown that this experimental procedure does not affect the observed genetic diversity (Böndel *et al.*, 2015). The populations cluster by geographic and genetic proximity and we denote therefore four groups: central (LA 1958, LA1963, LA2755, LA2931 and LA3111), northern (LA3784 and LA3786), southern coast (LA2750, LA2932 and LA4107), and southern mountain (LA4117(A), LA4118 and LA4330). A geographic map with the locations of all sampled populations is found in Figure 2.

**S2: 14 POPULATIONS NLR ENRICHMENT SEQUENCING**

**Probe design for NLR and CT genes.**

Probes for the NLRs were designed using the same target sequences as reported in (Jupe *et al.*, 2013)**.** In addition we used data from wild tomato NLR and a set of 14 CT genes, as described in (Stam *et al.*, 2016). The CT genes form a well describe set of 14 genes that have been used in previous studies (Städler *et al.*, 2008; Böndel *et al.*, 2015; Stam *et al.*, 2016) and have Sanger sequence data available. The main reason for inclusion of the CT genes is to validate the pooled sequencing approach and to verify the sensitivity and specificity of the SNP calling in pooled data sets. Exact functions of all the CT loci are not known. Böndel et al. (Böndel *et al.*, 2015) confirm that these CT loci evolve under neutrality and/or under purifying selection, thus these are suitable for the general inference of the species demography. Note that these loci are thus expected to show lower variation than the NLRs.

**Pooled enrichment sequencing**

Leaf material was collected from 10 mature plants per population and genomic DNA was extracted and quantified using a Qubit fluorometer (Life technologies). The samples were diluted and pooled per population to obtain 3 ng of high quality DNA with equal amounts per individual. The total DNA was prepared in 130 µl for gene enrichment. A sequencing library of enriched DNA was prepared using Agilents SureSelect XT with Custom Probes as in (Stam *et al.*, 2016). In short: DNA was sheared on a Covaris S220 to 600-800 bp. Size selection and cleaning was done using AMPure XP beads (Beckman Coultier) in two steps using 1.9:1 and 3.6:2 fragment DNA to beads ratio. The quality was assessed using a Bioanalyzer 2100 (Agilent). End repair, adenylation and adaptor ligation were preformed as described by Agilent. Pre-capture amplification was done using Q5 high fidelity PCR mixes. The amplified library was again quality checked on a Bioanalyzer 2100.

Hybridisation was performed as suggested for libraries <3 Mb. The libraries were indexed with 8bp index primers A01-B08 using Q5 PCR mix, quality was assessed using the Bioanalyzer and individual samples were quantified using Qubit. Eight samples were pooled in equal DNA amounts and the resulting pool was quantified by QPCR using the NGSLibrary quantification kit for Illumina (Quanta biosciences) and diluted down to a final concentration of 20 nM. Sequencing was done on an Illumina MiSeq sequencer to obtain 250 bp paired end reads. Sequencing was done with ZIEL - Institute for Food & Health, Core Facility Microbiome/NGS. Our probe set was based on predicted NLR sequences from *S. tuberosum, S. lycopersicum, S. pennellii,* known cloned NLR as described in (Jupe *et al.*, 2013) and NLR from *A. thaliana*. Previous studies have shown that similar probe sets successfully capture all NLR in closely related studies (Andolfo *et al.*, 2014).

**Mapping and Population analysis**

Read mapping and SNP calling were done using the same methods as described in Stam et al. (Stam *et al.*, 2016), with minor modifications. We used bedtools to extract read coverage data and calculate the depth per covered fraction of the targeted region in each sample for CT and NLR genes separately. SNPs were called using two callers GATK (McKenna *et al.*, 2010) and Popoolation (Kofler *et al.*, 2011). To verify the stringency of the filters and the cut-off values used in both programs, we compared the merged SNP calls to Sanger sequence data for three genes for all ten plants for several populations. After comparison, cut-offs were adjusted to obtain the best true SNP calls and both callers were run again. To obtain high quality data we assessed mapped reads for all genes for the populations. We then defined a subset of genes with uneven coverage and some instances or for which individual reads introduced large numbers of low frequency SNPs. These reads often overlapped with reads with missing mate pairs, suggesting mis-mapping likely due to insertion or deletion events or partial gene duplications. To avoid false positive genes (i.e. an over-estimation of the NLR diversity) these genes were removed from our analysis. Hence we identified a subset of 91 genes with high confidence annotations as well as high quality mapping data; even coverage, average depth > 30 and no gaps, in the majority of populations. The results in the main text are presented for this high quality subset only. Summary statistics $\pi$, $\pi_N$ and $\pi_S$ were calculated per site and gene in each population using a method based on the one of Nei and Gojobori (Nei & Gojobori, 1986) as implemented in SNPGenie (Nelson & Hughes, 2015; Nelson *et al.*, 2015).

**Population genetics and diversity assessment**

Per site values obtained for $\pi$, $\theta$ by SNPGenie were summarised and per population statistics were calculated. Per gene $\pi$, $\theta$, $\pi_N$ and $\pi_S$ statistics, were calculated over the HQ mapped and base-called gene length only to avoid bias due to missing data. Population averages were obtained by averaging per gene statistics. Further analyses were performed using R (R Core Team, 2015). We extracted per gene summary statistics and derived individual statistics per NLR class and genetic groups. $F_{ST}$ values were calculated for pairs of populations using the Hudson et al. estimator (Hudson *et al.*, 1992): $F_{ST} = (\pi_{between} - \pi_{within}) / \pi_{between}$. We define $\pi_{within}$ as the average $\pi$ for two populations, and $\pi_{between}$ as the nucleotide diversity called on the two populations with all HQ reads merged together. To obtain $\pi_{between}$ the whole pipeline was repeated for all of the 91 possible pairwise comparisons. (Bakker *et al.*, 2006) Results were analysed and visualised in R using the reshape (Wickham & Hadley, 2007) and ggplot2 (Wickham, 2009) packages. Maps were drawn using maps (Becker *et al.*, 2016) .

Statistical comparisons of groups or classes were done using analysis of variance as implemented in the aov() function. Where applicable post-hoc pairwise differences were calculated using tukeyHSD() function to account for multiple testing. All comparisons are reported as significantly different when $p < 10^{-5}$, unless stated otherwise.

As the SFS of some groups appear bias towards intermediate frequency variants we use conservatively the nucleotide diversity π and the derived $F_{ST}$ measure which are the least biased for allele frequency. Such inflation of intermediate frequency variants can be due to past demographic events but also an effect of our pooling procedure. As shown by the comparison to the previous results of Böndel et al., these measures are accurate in our datasets for the 14 CT loci (S figure 2D)

**S3: THREE GENOME COMPARISON / DEMOGRAPHY**

**Demographic inference with MSMC and simulation of summary statistics**

*Sequence data and heterozygosity*

We mapped the sequenced reads of the 3 sequenced plants from central (LA3111), southern mountain (LA4330), and southern coast (LA2932) populations against our *S. chilense* reference genome using BWA (mem, call -M with default parameters). SNPCalling was done using samtools (mpileup -q 20 -Q 20 -C 50). We restrict the variant calling to the 200 largest scaffolds of the *S. chilense* reference genome, comprising ~79.6Mb of sequence (mean length=398Kb, min=294Kb, max=1.12Mb). For LA3111 we found a total of 171,989 heterozygote sites (per scaffold mean: 859.94; sd: 921.88), 169,990 for LA4330 (mean: 849.95; sd: 894.62), and 176,630 for LA2932 (mean=883.15, sd=908.62).

*Demographic inference*

We used the multiple sequentially Markovian coalescent (MSMC v2) approach (Schiffels & Durbin, 2014) on unphased called genotypes to estimate both the effective population sizes and cross-coalescence rates over time using one individual per population representing each of the three main regions (central LA3111, southern mountain LA4330 and southern coast LA2932). The selected genomic regions (200 largest scaffolds of the *S. chilense* reference genome, each one of size longer than 200kb) showed 26X mean coverage for LA3111, 20X for LA4330 and 23X for LA2932. As recommended in the MSMC documentation, in addition to the variant matrix, we generate for each scaffold one mask file in bed format which restricts the analysis to the adequate covered regions of a given individual genome. In addition, we generate a mappability mask using Heng Li's SNPable tool (http://lh3lh3.users.sourceforge.net/snpable.shtml) which masks out all regions on the scaffolds on which short sequences cannot be uniquely mapped. Briefly, we used the SNPable tool set to divide the reference genome (200 largest scaffolds in our case) into overlapping short sequences of 100bp and align it back to the genome using BWA (aln -R 1000000 -O 3 -E 3). Then, bed files for the masked regions were generated with a python script (https://gist.github.com/danielecook/cfaa5c359d99bcad3200) and used in MSMC analysis as negative-mask. For the MSMC analysis we set the time segment patterning parameter (-p) to 15*1. For each run of the cross-coalescence analysis we used four haplotypes, two from each individual from the three populations using the -P 0,0,1,1. We plot the results assuming a mutation rate of $5\times10^{-8}$ per generation per base pair, and a generation time of 5 years (Städler *et al.*, 2008; Tellier *et al.*, 2011b).

We generate a visual confidence interval of the MSMC curves through replicates of the analysis with subsets of the dataset. This is performed as 10 independent runs of MSMC, each one run with a random subset of 100 scaffolds (out of 200) using the same settings as for the complete data set.

*Power analysis of the inference*

The MSMC method results is sensitive to the size of the genomic regions used for the analysis. Therefore, we test the ability of MSMC to recover the demographic history of simulated data sets with the same features as our empirical data for four specified demographic scenarios. We used the coalescent simulator ms (Hudson, 2002) to simulate independent scenarios of 1) different effective population size for each individual genome, and 2) two scenarios for the divergence and $N_e$ changes through time of the three genomes. For each scenario we implement 200 independent simulations (one per each reference genome scaffold) using the same length size of each scaffold (l) of our data set as well as the Theta (θ) and Rho (ρ) values estimated by MSMC. Effective population size and divergence time parameters for the simulations were based on the results obtained from the MSMC inference and previous knowledge (Böndel *et al.*, 2015).

We simulated two sets of single population scenarios:

1) Independent simulations for each population (central LA3111, southern mountain LA4330, and southern coast LA2932; S Figure 13-17) with constant effective population size (schematic model in S Figure 5B). The $N_0$ parameter was fixed to the highest value estimated by MSMC and adjusted by a coefficient to determine the population size of each population (i.e. 10* $N_e$ for central individual, 3* $N_e$ for mountain individual, and 10* $N_e$ for the coast individual). The ms command lines for each model were :

central: ms 2 1 -t 10*ϴ -r ρ l

southern mountain: ms 2 1 -t 3*ϴ -r ρ l

southern coast: ms 2 1 -t 2*ϴ -r ρ l

2) Independent simulations for each population (central, mountain and coast) with a past population expansion event (schematic model in S Figure 5C). Effective population sizes were set taking into account the estimated values by MSMC for each individual genome. We set the current effective population sizes as estimated by MSMC for the central population (0.59*locuslength), the mountain (0.83*locuslength) and the coast (0.83*locuslength). We set past population size expansion events for central to 10*e$N_0$ , 4*e$N_0$ for the mountain, and 2.5*e$N_0$ for the coast at times 0.12, 0.07, and 0.12 (generations/4$N_0$), respectively. Then a reduction to 3*e$N_0$ , 2*e$N_0$ for the mountain, and e$N_0$ for the coast, at time 0.39, 0.37, and 0.59, respectively. The ms command lines for each model model were:

central: ms 2 1 -t 10*ϴ -r ρ l -eN 0.39 0.1

mountain: ms 2 1 -t 3*ϴ -r ρ l -eN 0.37 0.33

coast: ms 2 1 -t 2*ϴ -r ρ l -eN 0.59 0.5

We also simulated two scenarios that include different divergence times and population size change parameters for the three populations, taken from the MSMC results for the empirical data as well as previous estimations for *S. chilense* (schematic model in S Figure 5D, E, (Böndel *et al.*, 2015)).

3) For the first scenario we set $N_0$=0.59*locus length (i.e. central population e$N_e$). We set the same current effective population for central, mountain and coast equal to $N_0$. We set past population size expansion events for to 10*e$N_0$ for central, 4*e$N_0$ for the mountain, and 2.5*e$N_0$ for the coast at times 0.12, 0.07, and 0.12 (generations/4$N_0$), respectively. Population split events were set to: mountain-central at time 0.25 and

coast-central at time 0.58. Finally, the ancestral population size was set to $N_0$ at time 0.57. For each population, a single diploid individual was sampled. The ms command to simulate the model was:

ms 6 1 -t ϴ -r ρ l -I 3 2 2 2
-en 0.11 2 4 -en 0.12 1 10 -ej 0.25 2 1
-en 0.12 3 2.5 -en 0.57 1 1
-ej 0.58 3 1

4) For the last scenario we also set $N_0$=0.59*locus length (i.e. central population e$N_e$) but the current effective population sizes were fixed to $N_0$ for central population, 2*$N_0$ for both mountain and coast populations. We set two past population size changes for central first to 16*e$N_0$ at time 0.11 and the second one to 10*e$N_0$ at time 0.57. We keep the mountain population size constant. For the coast population a single size change was set to 4*e$N_0$ at time 0.22. Population split events were set as: mountain-central at time 0.08 and coast-central at time 0.56. Finally, the ancestral population size was set to 2*$N_0$ at time 0.81. For each population, a single diploid individual was sampled. The ms command to simulate the model was:

ms 6 1 -t ϴ -r ρ l -I 3 2 2 2
-n 2 2 -n 3 2
-ej 0.08 2 1 -en 0.11 1 16
-en 0.22 3 4 -ej 0.56 3 1 -en 0.57 1 10
-en 0.81 1 2

All scenarios were simulated 10 times. Outputs of each simulation were converted to MSMC input using the ms2multihetsep.py script available in the MSMC-tools. Then, we run MSMC on these three simulated datasets with the same settings used for the empirical data and plotted the results compared with the empirical data (S Figure 5).

### S4: DEMOGRAPHIC INFERENCE USING APPROXIMATE BAYESIAN COMPUTATION

#### Demographic inference for all populations using ABC approach

We implemented a model-based demographic inference to obtain a neutral expectation of the genetic differentiation statistics used in this study ($F_{ST}$) under a well supported evolutionary scenario that represents the complete sampling (14 populations grouped in four regions). Thus, we used the Approximate Bayesian Computation (ABC) method (Beaumont *et al.*, 2002) to assess the support of three demographic models using a wide range prior parameter distributions for effective populations size ($N_e$), gene-flow and divergence times parameters (see table below for all parameters and prior distributions). Note that compared to the MSMC above, we have more populations and a more realistic population structure including gene flow between populations and groups. We assessed the support of three isolation-migration-island models: 1) independent divergence of the north, coastal and mountain groups from the central group, 2) same as the model 1 but the coastal group diverges from the highland group, and 3) same as the model 1 but the highland group diverges from the coastal group. Divergence processes were set to occur from three different unsampled (i.e. ghost) populations (see schematic figures of the models below). The gene-flow among populations from different groups was set as symmetric migration only from populations of the central group with the others (i.e. no migration between north-mountain, north-coast and mountain-coast populations).

Therefore, we have seven migration rates: three between populations of the different groups and four rates for within-group migration (each group is represented as an island model). Between-group gene-flow was constrained to be smaller than within-group gene-flow. Within a group, all populations were assumed to have the same population size for limiting the number of parameters in the model.

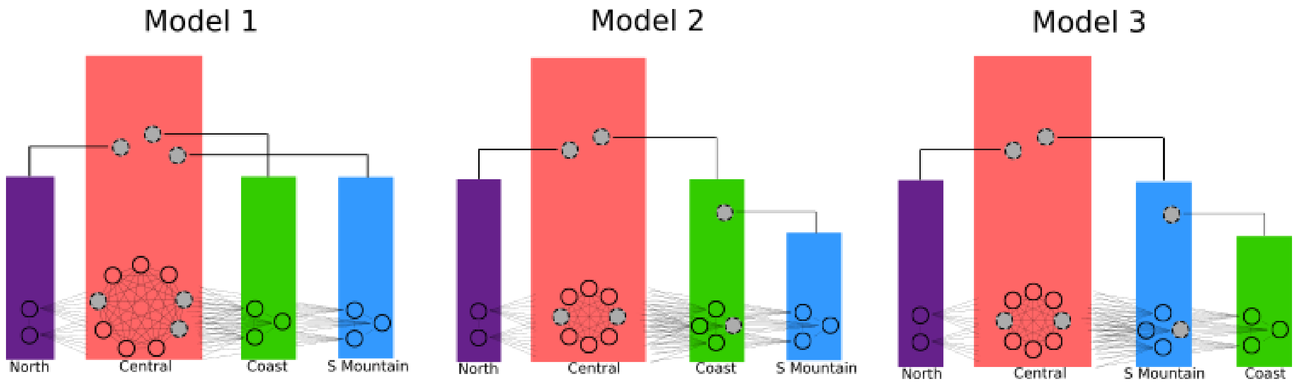


Figure1: Sketch of tested demographic models

Table 1: ABC parameters and observed distributions

| Parameter | Prior distribution (Min – Max) values | Posterior distribution Median (HPD 90% interval) |
|---|---|---|
| Effective population size central group populations | LogUniform (1 000 – 100 000) | 35 110 (25 689 – 44 767) |
| Effective population size north group populations | LogUniform (500 – 100 000) | 13 203 (3 171 – 29 100) |
| Effective population size coast group populations | LogUniform (500 – 100 000) | 8 042 (2 243 – 19 118) |
| Effective population size mountain group populations | LogUniform (500 – 100 000) | 16 451 (3 763 – 35 990) |
| Divergence time* north – central populations | LogUniform (100 – 2 000 000) | 12 607 (2 622 – 33 779) |
| Divergence time* coast – central populations | LogUniform (100 – 2 000 000) | 6 894 (735 – 20 158) |
| Divergence time* mountain – central populations | LogUniform (100 – 2 000 000) | 43 582 (3 878 – 108 851) |
| Migration within central group (MC) | logUniform (0.01 – 0.5) | 0.26 (0.18 - 0.36) |
| Migration within northern group | logUniform (0.01 – 0.5) | 0.08 (0.01 - 0.35) |
| Migration within coastal group | logUniform (0.01 – 0.5) | 0.1 (0.01 - 0.39) |
| Migration within mountain group | logUniform (0.01 – 0.5) | 0.08 (0.01 - 0.36) |
| Migration between central-northern populations | logUniform (0.01 – MC) | 0.07 (0.01 - 0.19) |
| Migration between central-coastal populations | logUniform (0.001 – MC) | 0.03 (0.01 - 0.07) |
| Migration between central-mountain populations | logUniform (0.001 – MC) | 0.1 (0.01 - 0.26) |
| Mutation rate (µ) | 1e-8 | - |
| Theta | $4*\mu*N_eC$ | 1.48 (1.09 - 1.89) |

* Divergence time in years ago (generation time 5 years).

For each model we simulated $10^6$ data sets using the software ms (Hudson, 2002) implemented with custom bash scripts. Prior parameter values were drawn from log-uniform distributions using the package KScorrect (Novack-Gottshall & Wang, 2018) in R. We used the sample_stats program included in the ms package and a custom PERL script written by N. Takebayashi (available at: http://raven.wrrb.uaf.edu/~ntakebay/teaching/programming/coalsim/scripts/msSS.pl) to calculate proportion of polymorphic sites within each population ($\pi_S$ within), and between populations ($\pi_S$ between). Note that because we want to test the neutral distribution, we restrict to the calculation of the observed summary statistics to synonymous sites only from our complete CT and NLR data set. Pairwise $F_{ST}$ values were calculated using the Hudson et al. Estimator ($F_{ST} = (\pi_{between\ -}\ \pi_{within})/\ \pi_{between}$; Hudson *et al.*, 1992). The scripts used to draw parameter values from prior distributions, coalescent simulations for each model and calculation of summary statictics can be found in https://github.com/gsilvaarias/NLR-ABC. For initial inspection of the simulated and observed summary statistics, we implemented a PCA for a randomly-taken $10^4$ simulated summary statistics of each model using the prcomp function in R (see figure below). We calculated the posterior probabilities for each demographic model using the ABC package (Csilléry *et al.*, 2012) in R. We assessed two model selection methods: the multinomial logistic regression method and the neural network approach. Those methods use adjustment-based nonlinear approaches that minimize departures from linearity and homoscedasticity and allow a reduction in the dimensionality of the SuSt via internal projections on lower-dimensional subspaces (Blum & François, 2010). We used three threshold level of simulations retained to estimate the posterior probabilities of each model (0.1%, 1% and 10%).

To calculate posterior parameter distributions, we performed post-rejection adjustments within the prior parameter value bounds (Blum & François, 2010) using the neural network approach available in the abc package (Csilléry *et al.*, 2012) in R. A posterior predictive verification taking the 10,000 best simulations was used to draw a new posterior parameter distribution. We plotted the $F_{ST}$ values (S Figure 6B) and histograms of the calculated summary statistics of the posterior simulations (S Figure 6C) to assess the fit with the observed values. Finally, we implemented a posterior PCA for $10^4$ summary statistics randomly taken from simulations of the tested models, those from the posterior parameter distribution, and the observed data (S Figure 6A).

**ABC Model selection and parameter estimation, results**

Rejection analyses show the model of independent divergence of the north, coast and mountain groups from the central group (model 1) as the the best supported model given our genetic data set . Posterior parameter estimations were obtained from 0.1% of the prior samples (10000 out of $10^6$ simulations) of the best supported model with post-rejection adjustments using a neural network algorithm. Posterior estimations for the effective population sizes show higher $N_e$ for populations from central group than for the others. Populations from the coastal group showed very low $N_e$ estimation. In contrast to the MSMC results, the divergence time for the coast populations was estimated to be more recent than divergence of the mountain populations. Migration estimates show high values for population within the central group. The other groups show very low migration estimates and similar to the migration rates among groups (all of them have zero for lower HPD bounds). The lowest estimates for migrations was detected among central and coast populations, while gene-flow between central and mountain populations was found to be markedly higher and similar to the with groups estimates. Posterior predictive verification show that simulations using posterior parameter distributions are able to recover the observed summary statistics values fairly well as seen in the PCA scatterplot and in the summary statistics distribution plots (S Figure 6).

Table 2: Posterior probability for each model in the ABC based on the number of simulations retained.

| **Rejection method** | **Model** | **Threshold (number of simulations retained closer to observed data)** | | |
|---|---|---|---|---|
| | | **3 000** | **30 000** | **300 000** |
| Multinomial logistic regression | 1 | 0.63 | 0.97 | 0.97 |
| | 2 | 0 | 0 | 0 |
| | 3 | 0.37 | 0.03 | 0.03 |
| Neural networks | 1 | 0.37 | 0.99 | 0.94 |
| | 2 | 0.05 | 0 | 0.03 |
| | 3 | 0.58 | 0.01 | 0.03 |

### S5: SIMULATION OF NEUTRAL (GENOMIC) $F_{ST}$ RANGES

**Calculation of neutral interval of pairwise $F_{ST}$ measures for the identification of outlier NLR based on MSMC and ABC results**

The two demographic scenarios inferred from MSMC analysis that considered divergence and past $N_e$ changes described above were used to create neutral 'null' distributions of the pairwise population $F_{ST}$ values. For each scenario we performed 10,000 simulations using the mean size length of the R-genes assessed in this study (mean length = 2149 bp). We used ms with to simulate 10 diploid samples per population

Model 1:

ms 60 10000 -t 7.7 -r 0.94 2149 -I 3 20 20 20
-en 0.11 2 4 -en 0.12 1 10 -ej 0.25 2 1
-en 0.12 3 2.5 -en 0.57 1 1
-ej 0.58 3 1

Model 2:

ms 60 10000 -t 7.7 -r 0.94 2149 -I 3 20 20 20
-n 2 2 -n 3 2
-ej 0.08 2 1 -en 0.11 1 16
-en 0.22 3 4 -ej 0.56 3 1 -en 0.57 1 10
-en 0.81 1 2

$F_{ST}$ values were calculated using the Hudson et al. estimator (Hudson *et al.*, 1992) as obtained for the ABC analysis (see above).

To detect the outlier NLR, the highest $F_{ST}$ values in each of the three pairwise comparisons were selected as a cut-off. Similary, from the ABC posterior parameter estimations we generated a set of neutral distributions of $F_{ST}$ values for all population pairwise comparisons (14 populations) based on 30,000 loci defined by the average length of our NLRs (2149) and genomic population recombination rate estimated with MSMC

($4.5x10^{-9}$ – $1.1x10^{-8}$ per site per generation). We emphasise here that this cut-off is very stringent and conservative (Figure 6B).

**Expected distribution of pairwise $F_{ST}$ measures under selective scenarios**

We performed forward genetic simulations using the software SLiM 3.2 (Haller & Messer, 2017) to explore the values and distributions of  pairwise $F_{ST}$ due to selection during to the southward colonization process in *S. chilense.* The demographic model is based on  the demographical parameters inferred from MSMC results (Fig 5; SFig 5D). We simulated a linear genome of 9.1 Mb made up of 100 repetitions of 91 loci of 1 kb. Each of these regions consisted of neutral mutations with 10 scattered sets of 10bp stretches which have mutations experiencing either 1) strong positive (selection coefficient *s*=0.01), 2) balancing (modelled as negative frequency-dependent) selection . Selection occurs only in the  southern populations and the central group has only neutral evolution. For computational efficiency, we scaled the event times and population sizes by a factor of 0.1. We set the per-site mutation and recombination rate to $1\times10^{-7}$. This ensures that there is some amount of linkage disequilibrium (as expected in real data) but the 91 loci do evolve independently from one another.  For each replicate we obtained an output in vcf format a sample of 60, 30 and 30 diploid individuals for central, mountain and coast populations, respectively, and calculated Weir and Cockerham $F_{ST}$ for neutral sites and the sites under selection using vcftools (Danecek *et al.*, 2011).

**S6 OUTLIER NLR**

**Definition of outlier NLR and classification**

The simulated $F_{ST}$  values were imported into R. For each pairwise comparison between the populations, we selected the NLRs that fell outside the maximum simulated value (out of 30,000 simulations). Again, we highlight here the stringency of this criterion. The data were transformed using the packages reshape (Wickham & Hadley, 2007) and plyer (Wickham, 2011) and visualised using the ggplot2 package (Wickham, 2009).

Main habitat adaptation genes were defined by selecting the genes that occur as outlier in at least one third of the possible pairwise population comparisons between two groups. For example, *g*. when comparing the central (6) vs coastal (3) populations, a gene is qualified as a main habitat gene if it is found  as an outlier more than 6 times out of the 18 possible pairwise population comparisons. This criterion insures selection of those NLRs that are outliers in comparisons involving at least two out of the three derived populations (for the southern groups), which is a strong indication of fixation and/or selection of common alleles in several populations of the same group. All genes that do not match this criterion (e.g. appear in fewer comparisons or only in comparisons within a geographic group) were defined as local adaptation “fine-tuning” genes. These do not show a consistent, geography-based evolutionary pattern. The NLR clades associated to the main habitat adaptation and local adaptation NLRs were again visualised using ggplot2.

To test whether the relative abundance of the NLR classes in the main and local adaptation lists  did arise by chance, we randomised the $F_{ST}$ values 1) within the whole data set, and 2) within the total set of selected outlier NLRs, and subsequently reran our analyses. Using these randomisation outputs  we calculated the average number of major genes that can be identified (under 1000 whole dataset randomisation) or the mean number of NRC genes that are classified as major gene (in 1000 randomisations following procedure 2 within outliers). These averages are interpreted as the expectations for the two quantities. We estimated the confidence interval from the random sampling (mean ±2σ). The observed number of major genes we find (17) is much larger than the expectation which has mean 3.5 (and C.I. 0.1 – 7.2]). Similarly, the observed fraction of NRC genes amongst the main habitat adaptation genes is 5 and larger then the expected 1 NRC (CI. [0.6-2.3]).

**S7 DATA AVAILABILITY**

The *S. chilense* enrichment sequencing data are available on NCBI in BioProject PRJNA474589,
